# CD44v6 Drives Tumor Aggressiveness and Chemoresistance in Bladder Cancer

**DOI:** 10.1101/2025.08.27.672563

**Authors:** Iris Lodewijk, Carolina Rubio, Pontus Eriksson, Ignacio A. Reina, Esther Montesinos, Miguel Alonso-Sánchez, Laura García-Gómez, Álvaro Martín de Bernardo, Lucía Morales, Cristian Suárez-Cabrera, Omaira Alberquilla, Rebeca Sánchez-Domínguez, Félix Guerrero-Ramos, Rosa García-Martin, Lucía Parrilla, José L. Rodríguez-Peralto, Daniel Castellano, Jesús M. Paramio, Gottfrid Sjödahl, Marta Dueñas

## Abstract

Bladder cancer is a fast-moving and recurrent malignancy where survival hinges on early detection and precise risk stratification. The search for robust biomarkers is urgent, and CD44v6 has emerged as a compelling candidate. In this study, we reveal that CD44v6 is not merely a marker but a driver of urothelial carcinoma aggressiveness. Through integrated clinical and functional analyses, we show that high CD44v6 expression is strongly associated with poor patient outcomes. Mechanistic investigations demonstrate that CD44v6 amplifies the proliferative, migratory, and invasive potential of bladder cancer cells, while conferring marked resistance to cisplatin. These findings position CD44v6 at the intersection of tumor progression and therapeutic failure, underscoring its value as both a prognostic biomarker and a promising therapeutic target. Targeting CD44v6 could pave the way for strategies that curb disease aggressiveness and overcome chemoresistance in bladder cancer.

## 1. INTRODUCTION

Bladder cancer (BC) is a highly prevalent urologic malignancy that comprises about 5% of all cancer cases and represents the 10^th^ most diagnosed cancer worldwide ^1^. It has become an important global health problem and a major cause of morbidity and mortality, accounting for approximately 615,000 new cases and 220,000 BC deaths in 2022 alone (Bray et al., 2024). While early detection and treatment improve patient outcomes, the challenges associated with disease recurrence and progression persist. At diagnosis, roughly 75% of cases fall within the non-muscle-invasive BC (NMIBC) category; the remaining 25% are classified as muscle-invasive BC (MIBC) ^3^. The current paradigm of patient management is intricately linked to this pathological classification, with NMIBC being typically addressed through bladder-sparing approaches, specifically transurethral resection with adjuvant intravesical therapies ^4^. In contrast, MIBC necessitates an aggressive approach involving radical cystectomy, lymphadenectomy, and platinum-based chemotherapy administered in adjuvant or neoadjuvant contexts, although a number of patients are candidates to aggressive bladder-sparing approaches such as trimodal therapy ^4^. In a notable proportion of MIBC cases the disease advances to the metastatic stage (mBC), an incurable condition of extremely low overall survival. Traditionally, platinum-based combination chemotherapy has served as the cornerstone of systemic treatment for patients with mBC ^5^. Recent advancements have positioned immune checkpoint inhibitors (ICI) as a promising therapeutic avenue, in some cases demonstrating profound and enduring responses in mBC patients who have progressed after platinum-based chemotherapy ^6–10^.

There are other novel therapeutic alternatives that have further transformed the landscape of mBC therapy. These include antibody-drug conjugates (ADC), FGFR inhibitors tailored for tumors harboring FGFR alterations, and more recently, Enfortumab vedotin (EV) ^11^ and Sacituzumab govitecan in patients with metastatic urothelial carcinoma ^12^. However, it is noteworthy that response rates in patient cohorts remain relatively modest, and sometimes metastatic lesions acquire resistance to these interventions due to downregulation of specific molecular targets ^13^. Therefore, a need exists for the identification and evaluation of a larger number of tumor-associated antigens (TAAs), in order to devise more precise treatment strategies comprising a battery of drugs targeting all tumor types associated with the individual characteristics of each tumor in each treatment period.

Aberrant expression and dysregulation of CD44, a transmembrane glycoprotein that participates in multiple physiological processes, has been shown to contribute to cancer initiation and progression ^14^. CD44 promotes epithelial-mesenchymal transition (EMT) and is a common biomarker of cancer stem cells (CSCs) ^15^. It exists as multiple variants, including the standard CD44s and several other isoforms ^16^. Among the latter, CD44v6 is overexpressed in several cancers, including head and neck squamous cell carcinoma (HNSCC), ovarian, colorectal, and thyroid cancer ^17–19^. In addition, CD44v6 has been shown to be involved in cell invasion and metastasis ^18,20,21^ and is therefore considered a potential therapeutic target for cancer; as a matter of fact, targeting CD44v6 for radiotherapy has yielded promising results in preclinical and clinical trials ^22^.

In BC, CD44v6 is also considered a promising diagnostic and prognostic biomarker due to the fact that it is expressed on BC cells but not on normal bladder cells (Omran & Ata, 2012 and Lodewijk et al 2023). Unfortunately, in contrast with the abundant data evidencing the utility of CD44v6 as a potential prognostic factor in cancers such as HNSCC, no studies have focused on the molecular mechanisms underlying its tumorigenicity within the context of BC. Since an understanding of the biological behavior underlying CD44v6 expression if fundamental to refine its use in clinical practice and explore innovative therapeutic approaches that can ultimately improve patient outcomes, the present study is aimed at comprehensively investigating the molecular characteristics of CD44v6-expressing BC, focusing on the diagnostic, prognostic, and therapeutic value of this protein. We examine whether CD44v6 expression can be used as a sensitive and specific biomarker for BC detection and investigate if there is a positive correlation between its expression levels and tumor grade, stage, and recurrence risk. Additionally, we propose that targeting CD44v6 may enhance the efficacy of existing therapies and provide new therapeutic avenues for BC treatment.

## 2. MATERIALS & METHODS

### 2.1 The Cancer Genome Atlas (TCGA) data analyses

We used the web-based tool TSVdb ^24^ (http://www.tsvdb.com/plot.html) to access TCGA, a comprehensive public resource providing a wide array of genomic data across various cancer types, to download Level_3 RNA sequencing (RNAseq) gene expression data for Bladder Urothelial Carcinoma (BLCA). The selected gene was *CD44*, and *CD44v6* gene expression data were obtained from alternative splicing isoform_uc001mwc. Accession took place in June, 2024 and the resulting RNA-Seq by Expectation-Maximization (RSEM) normalized data, corresponding to samples from 406 BC patients, were transformed by log2. These data were used for survival analysis as well as gene expression comparisons across different disease stages and sex, according TCGA classification (Supplementary Data 1). Data validation was performed using the “Lund s265 cohort”, dataset deposited in: https://zenodo.org/records/10362517 ^25^. CD44 isoform expression was quantified using the NF-core pipeline, employing Spliced Transcripts Alignment to a Reference (STAR) for alignment and Salmon for transcript-level quantification.

### 2.2 Patient series

Patient enrollment and data collection procedures were supervised and approved by the Ethical Committee for Clinical Research of the 12 de Octubre university hospital. The study used samples from 46 NMIBC and 42 MIBC patients (Supplementary Table 1). The samples, as well as clinical united data of the patients, were provided by Biobanco i+12 in Hospital 12 de Octubre in coordination with the Spanish Hospital Biobanks Network (RetBioH; http://www.redbiobancos.es), following standard operating procedures after approval by the Ethical and Scientific Committee (CEIC16-011). Written informed consent was obtained from all patients/participants prior to the study.

### 2.3 Tissue microarrays and Immunohistochemistry (IHC)

Initial sections were cut, stained with hematoxylin/eosin (HE) and evaluated by an experienced pathologist to verify previous histopathological findings and identify representative areas for tissue microarray (TMA) construction. Two tissue cores (diameter 0.6 mm and thickness of 4 μm) per patient from the selected tumor area were then used to construct TMAs with a manual tissue arrayer (Beecher Instruments, Sun Prairie, WI). The TMAs were stained with HE and examined to confirm the presence of representative tumor tissue (at least 70% of tumor cells).

Sections of formalin-fixed paraffin-embedded (FFPE) tumors were deparaffinized and blocked with hydrogen peroxide (0.3%) in methanol. Antigen retrieval was performed in sodium citrate buffer (citric acid monohydrate 1.8 mM, trisodium citrate dihydrate 8.2 mM, pH=6.0), using a pressure cooker (Dako Agilent Technologies). The sections were then blocked with 10% horse serum (HS) for 30 min. at 37°C, followed by overnight incubation with the corresponding primary antibody (Supplementary Table 2) diluted in 10% HS. Next, the samples were incubated for 1 hour at room temperature with biotin-conjugated secondary antibodies (Supplementary Table 2) diluted in HS 10%, followed by incubation with avidin-peroxidase (VECTASTAIN®Elite® ABC Kit; Vector Laboratories) to amplify the signal. The slides were developed with diaminobenzidine as a substrate (DAB Substrate Kit; Vector Laboratories), following manufacturer’s instructions. Positive control slides were included to confirm antibody reactivity, and the sections were counter-stained with hematoxylin.

CD44s and CD44v6 expression was scored based on quantity and intensity of the staining (score 0: negative samples; score 3: complete and very intensive staining). Scoring was performed by two independent investigators, reconciling discordant results by joint examination and discussion. Internal controls for reactivity of each antibody and tissue controls for the series were added according to previously published methods.

### 2.4 Immunofluorescence

Tissue sections were deparaffinized and antigen retrieval was performed as previously described (Section 2.3). After incubation with NH_4_Cl 100 mM for 30 min. at room temperature (RT) to quench background fluorescence, the sections were blocked with 10% HS for 30 min. at 37°C, followed by incubation with primary antibodies (Supplementary Table 2) diluted in 10% HS. Next, the sections were washed and incubated with secondary antibodies (Supplementary Table 2) diluted in 10% HS. After further washing, the coverslips were inverted onto mowiol (Fluka) mounting medium with 4’,6-diamidino-2-phenylindole (DAPI) (Dilution 1:1000; Vector Laboratories) on microscope slides, viewed with a Zeiss digital microscope and photographed.

### 2.5 Cell lines and reagents

ATCC Human BC cell lines SW-780, MGH-U3, MGH-U4, RT-4, 5637, RT-112, T24, UM-UC-3, J82, and 253J of known genomic characteristics were initially evaluated (Supplementary Table 3) ^26^. 5637 cells were grown in RPMI 1640 medium (Gibco-Thermo Fisher Scientific Inc.) and the remaining cell lines were maintained in DMEM medium (Gibco-Thermo Fisher Scientific Inc.), supplemented with 10% fetal bovine serum (FBS) (Hyclone) and 1% Antibiotic/Antimycotic (A/A) (Gibco-Thermo Fisher Scientific Inc.) at 37°C in a humidified atmosphere of 5% CO_2_. All cell lines were authenticated by short tandem repeat allele profiling and frequently tested for Mycoplasma using Venor®GeM OneStep (Minerva Biolabs) following manufacturer recommendations.

### 2.6 Flow cytometry & Fluorescence-Activated Cell Sorting

*Receptor expression* - Fc receptors were blocked with 0.2 μl/100,000 cells of FcR Blocking Reagent (Miltenyi Biotec) (final volume 100 μl) for 10 min. at 4°C, after which the cells were stained with primary antibodies (Supplementary Table 2) in phosphate-buffered saline (PBS) (Gibco-Thermo Fisher Scientific Inc.) for 30 min. in the dark at 4°C. Unstained samples were included as negative controls. The samples were centrifuged (5 min., 1400 rpm, RT) and resuspended in PBS containing 2 μg/ml DAPI (Roche) for final analysis. CD44 and CD44v6 receptor expression data were acquired using an LSRFortessa™ cell analyzer (BD Biosciences) and BD FACSDiva software (BD Biosciences). CD44 and CD44v6 receptor expression was analyzed using FlowJo™ v10.7 software (Becton, Dickinson and Company). Median Fluorescence Intensity (MFI) was calculated using FlowJo™ v10.7 software, subtracting the MFI of negative controls. Each experiment was performed at least 3 times.

*Cell sorting* - J82 and RT-112 cells were stained with FITC anti-human CD44 antibody and PE anti-human CD44v6 antibody as previously described (Supplementary Table 2). 1-3% of cells with highest CD44, lowest CD44 and highest CD44v6 receptor expression were isolated from each cell line using the BD FACS InfluxTM Cell sorter (BD Biosciences) (Supplementary Figure 1; Figure 3C). Each experiment was performed 3 times.

*Xenograft-derived tumors* – Cell suspensions derived from disaggregated tumors were stained with Zombie Aqua^TM^ dye (Dilution 1:150 for 1x10^6^ cells) (Biolegend) for 20 min. at 4°C. Fc receptors were blocked, and cells were stained according to the abovementioned protocol, using antibodies against CD44, CD44v6, CD326 (EpCAM) and CD45 (Supplementary Table 2). Fluorescence Minus One (FMO) controls were included in this experiment, as well as unstained samples representing negative controls. Finally, the samples were centrifuged (5 min., 1400 rpm, RT) and resuspended in PBS for analysis. The LSRFortessa™ cell analyzer and BD FACSDiva software were used for data acquisition, analyzing the resulting data in the OMIQ Data Science Platform (Dotmatics).

### 2.7 RNA extraction

Total RNA from cell lines and FFPE tissue sections was isolated with the miRNeasy Mini Kit (QIAGEN) and miRNeasy FFPE Kit (QIAGEN), respectively, following instructions from the manufacturer. A final DNAse treatment step (RNAse-Free Dnase Set; QIAGEN) was included in both cases. RNA concentration was measured with a Qubit 4 Fluorometer (Thermo Fisher Scientific), using either the Qubit RNA broad range assay kit (Thermo Fisher Scientific) or the Qubit RNA high sensitivity assay kit (Thermo Fisher Scientific). RNA samples were stored at -80°C.

### 2.8 RNA sequencing

Three independent biological replicates of total RNA (≥500 ng) from each cell line (from both RT-112 and J82: parental, CD44 Low, CD44 High and CD44v6) were used for RNAseq analysis. All submitted samples had an RNA integrity number (RIN) higher than 9.4, as determined in an Agilent 2100 Bioanalyzer (Agilent Technologies). The samples were shipped to Macrogen RNA-seq services (Seoul, South Korea) for rRNA depletion with Ribo-Zero Gold (Illumina), library preparation with the SMARTer Stranded total RNA kit (TaKaRa) and analysis on the NOvaSeq 6000 platform, 150PE (150 x 2bp) (Illumina). A total of 40 M reads/sample (6 Gb/sample) were obtained.

In the case of FFPE patient tissue sections, total RNA samples (≥500 ng) were submitted for RNAseq analysis using the SureSelect RNA Direct Library (Illumina) in the NovaSeq 6000 platform, 100PE (100 x 2bp) (Illumina). All submitted samples had a DV200 (%) (percentage of RNA fragments > 200 nucleotides) higher than 25%, as measured with an Agilent 2100 Bioanalyzer (Agilent Technologies). A total of 60 M reads were obtained per sample (6 Gb/sample).

Differential expression analyses using the RNA-seq data were performed with the DESeq2 package (version 1.42.0) in R (version 4.3.2) ^27,28^. Normalized gene expression data were transformed into regularized log2 scale. Principal component analysis (PCA) was performed in R, visualizing the results via 3D scatter plots (https://tools.altiusinstitute.org/cubemaker/). Genes with a foldchange >1.5 for upregulated genes and <-1.5 for downregulated genes and an adjusted p-value <0.05 were selected for clustering and Enrichr analysis (https://maayanlab.cloud/Enrichr/). Hierarchical clustering was performed using Multi-Experiment Viewer (MeV 4.9.0) ^29^. Unsupervised hierarchical clustering was applied using Pearson correlation and average linkage. Heatmaps were represented using MeV version 4.9.0. Gene set enrichment analysis (GSEA) was performed using GSEA version 4.3.2 and the Molecular Signature Database (MSigDB) ^30,31^. The h.all.v7.5.symbols.gmt (Hallmarks) gene set database was used as the gene set collection analysis. GSEA was performed using 1000 permutations, with maximum and minimum sizes for gene sets of 500 and 15, respectively. Gene set variation analysis (GSVA) was performed with the GSVA package in R (v1.50.5) and a hallmark gene set ^32^. Data was represented by a heatmap using MeV version 4.9.0. The Datasets have been deposited in the Gene Expression Omnibus.

RNA-Seq FFPE samples were classified into molecular subtypes using three specialized tools. BLCAsubtyping was used to classify BC subtypes based on gene expression models, following the guidelines of Kamoun et al. (2020). The resulting data were visualized in the SRPLOT Bioinformatics platform (https://www.bioinformatics.com.cn/en) ^34^.

Analysis of miRNA expression in FFPE patient tissue sections was performed from total RNA samples (≥100 ng), using nCounter technology (NanoString Technologies). nCounter employs molecular barcodes, directly tagging RNA with a target-specific capture probe and a target-specific reporter probe containing a labeled barcode. Preparation and analyses were performed with the Human v3 miRNA Assay (NanoString Technologies), which includes 827 miRNAs and 5 mRNAs, following instructions from the manufacturer. Normalization was performed with nSolver software (Nanostring Technologies), correcting for the expression of technical controls and 25 internal reference controls included in the panel. Clustering was done using those miRNAs exhibiting a foldchange >1.5 in the case of upregulation or <-1.5 for downregulation and an adjusted p-value <0.05. Hierarchical clustering was performed using Multi-Experiment Viewer (MeV 4.9.0). Unsupervised hierarchical clustering was applied using Pearson correlation and average linkage. Heatmaps were represented using MeV version 4.9.0. Datasets have been deposited in the Gene Expression Omnibus.

### 2.9 Quantitative PCR with reverse transcription (RT–qPCR)

Reverse transcription was performed on 1 ug of total RNA from BC cell lines, 100 ng of total RNA from sorted cell populations and 30 ng of total RNA from xenograft-derived tumors using the High-Capacity cDNA Reverse Transcription Kit (Thermo Fisher Scientific). PCR was performed in a QuantStudio 5 Dx Real-Time PCR System (Thermo Fisher Scientific) with GoTaq PCR master mix (Promega) and unpurified cDNA as a template (1 μl of cell line-derived cDNA or 1.5 μl of xenograft-derived tumor cDNA). The oligonucleotides used are listed in Supplementary Table 4. Melting curves were performed to verify specificity and absence of primer dimerization. Reaction efficiency was calculated for each primer combination, and *TBP/GUSB* was used as reference gene for normalization.

### 2.10 Cell proliferation assay

*XTT assay* - RT-112 and J82 cells were seeded at a density of 3,500 and 1,500 cells/well, respectively, in a 96-well plate and incubated at 37°C in a humidified atmosphere of 5% CO_2_. Cell proliferation was assessed with the XTT Cell proliferation kit II (Roche), measuring the results in a GENios pro microplate reader (Tecan) at time points 0, 24, 48 and 72h (Supplementary Figure 2A). Time point 0 is actually absorbance at 16h post-seeding, and was used as reference for data normalization. Mean net absorbances (average of six replicates) were background corrected by subtracting the mean of medium-only blanks. Each experiment was performed at least 3 times.

### 2.11 Cell migration assay

*Wound Scratch Assay* - RT-112 and J82 cells were seeded at a density of 300,000 and 200,000 cells/well, respectively, in a 6-well plate and incubated at 37°C in a humidified atmosphere of 5% CO_2_. After 24 hours, the complete growth medium was removed, and cells were maintained in FBS-deprived DMEM medium (0% FBS) for 72 hours. A sterile 20-200 μl pipette tip was used to gently scratch a cross in each well. Scratch closure was monitored and imaged at time points 0, 24, 48 and 72 h using a Nikon Eclipse Ts2R microscope equipped with a Leica DFC 7000T camera (Leica) and the Leica LasX (Leica) software package (Supplementary Figure 3A). Time point 0 represents image acquisition directly after cross scratching, and was used as reference for data normalization. Scratch image analysis was performed with Fiji (ImageJ) ^35^. Each experiment was performed at least 3 times.

### 2.12 Cell invasion assay

*Boyden Chamber Assay* – Corning® BioCoat® Control Inserts (Corning) and Corning® BioCoat® Matrigel® Invasion Chambers (Corning) with 6.4 mm diameter Polyethylene terephthalate (PET) membrane filters of 8 µm pore size were used to form dual compartments in a 24-well tissue culture plate. RT-112 and J82 cells were seeded at a density of 100,000 and 50,000 cells/well, respectively, into the upper compartment of the dual chambers in supplemented DMEM medium. Supplemented DMEM medium was also placed in the lower compartment and plates were incubated at 37°C in a humidified atmosphere of 5% CO_2_. After cell attachment to the membrane, the supplemented DMEM medium of the upper compartment was replaced by FBS-deprived DMEM (0% FBS). After 24 hours, cells were detached from the membrane filter with Accutase®Cell Detachment Solution (BioLegend) and collected separately from the upper and lower compartments. After staining with CD44 and CD44v6 antibodies, the cells were then analyzed in a LSRFortessa™ cell analyzer with BD FACSDiva software, as previously described (Section 2.6). Precision count beads^TM^ (BioLegend) were added for quantification and normalization of traversed cells (Supplementary Figure 4A). The resulting data were analyzed using FlowJo™ v10.7 software (Becton, Dickinson and Company). Each experiment was performed in triplicate and repeated at least twice.

### 2.13 Platinum sensitivity assay

*XTT assay* - RT-112 and J82 cells were seeded at a density of 6,000 and 5,000 cells/well, respectively, in a 96-well plate and incubated at 37°C in a humidified atmosphere of 5% CO_2_. After 24 hours, the cells were treated for another 24 h with serial dilutions of cisplatin (Selleckchem) dissolved in PBS (1 mg/ml stock solution; 3.33 mM), ranging from 6 to 50 μM for RT-112 and from 6 to 400 μM for J82. Cell viability was assessed with the XTT Cell proliferation kit II (Roche), reading the data in a GENios pro microplate reader (Tecan) (Supplementary Figure 5). After subtracting medium-only background absorbance, the data (mean of six replicates per drug concentration point) were normalized as percent of vehicle control (PBS). The maximum mean inhibitory concentration (IC_50_) value of cisplatin for each cell line was determined using GraphPad Prism version 9 (GraphPad Software). Each experiment was performed at least 3 times.

### 2.14 Tumor xenografts

All animal experiments were conducted in compliance with CIEMAT guidelines and approved by the Animal Welfare Department of Comunidad de Madrid (Spain; PROEX 150.8/21). RT-112 cells were trypsinized and resuspended in a mixture (1:1) of PBS with Vitrogel (BD Biosciences). Five million cells in suspension were subcutaneously injected in each flank of 16 healthy immunocompromised nude (NMRI-FoxN1^nu/nu^) female, 8-10 weeks old mice (Janvier Saint-Berthevin, France). Tumor height and width were measured three times a week, using an electronic digital caliper (mm). Tumor volume was calculated as follows:

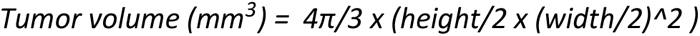

Tumor growth was monitored until tumor size became nearly unacceptable in the most affected individuals (humane endpoint). Tumor size and weight were measured after euthanasia.

### 2.15 Tumor disaggregation

Tumors were carefully sliced into 1-2 mm^3^ pieces by microdissection. The fragments were incubated in an enzymatic cocktail of DNase I (100 µg/ml) (Roche), collagenase P (200 µg/ml) (Roche) and dispase II (800 µg/ml) (Sigma-Aldrich) in DMEM medium at 37°C, vortexing every 20 min. for 3 to 4 times. Digested tissue samples were filtered through a 40 µm cell strainer, and the resulting cell suspensions were centrifuged (5 min., 1200 rpm, RT). The cells were resuspended in 1x BD Pharm Lyse™ lysing solution (BD Biosciences) and incubated for 10 min. at RT in the dark, followed by centrifugation (5 min., 1200 rpm, RT) and finally resuspension into PBS.

### 2.16 Statistical analyses

Comparisons using gene expression data from TCGA were made with unpaired t-tests. Contingency analyses were performed with Fisheŕs exact test or Chi-square test, as appropriate. Differential CD44 and CD44v6 receptor expression for each BC cell line was evaluated with unpaired t-tests. Maintenance of differential CD44 and CD44v6 receptor expression as well as invasion capacity among newly established cell lines were analyzed by a one-way ANOVA test. Differential percentages of either CD44+/CD44v6+ or CD44+/CD44v6-between seeded and migrated cells in the cell invasion assay were determined by unpaired t-tests. The statistical analysis of proliferation and migration capacity in newly generated cell lines was done with a two-way ANOVA test. *In vivo* tumor growth was evaluated by means of a two-way ANOVA test. Discrimination between tumor sample size or weight was made using the median. CD44 and CD44v6 receptor expression in the implanted tumors was analyzed with a one-way ANOVA test. Differences in cisplatin sensitivity between the cell lines were determined by the extra sum-of-squares F test, comparing IC_50_ values. GraphPad Prism 10.1.2 was used. In all cases, a p-value < 0.05 was considered significant.

## 3. RESULTS

### 3.1 CD44 and CD44v6 are expressed in the majority of urothelial tumors

In order to assess the relevance of CD44 and CD44v6 in BC, we retrieved the available *CD44* and *CD44v6* gene expression data corresponding to Bladder Urothelial Carcinoma (BLCA) at TCGA, which belonged to samples from 406 BC patients. Gene expression analysis of *CD44* data showed no clear association with epidemiological and clinical factors such as survival, disease stage or sex (Supplementary Figure 6A-6C). Likewise, we failed to find an association between *CD44v6* expression levels and overall survival or sex. However, *CD44v6* did exhibit significantly increased expression in samples from stage III and stage IV disease (Figure 1A; Supplementary Figure 6A & 6C). Additionally, the percentage of *CD44v6* positive samples increased from 0% in stage I disease samples to almost 60% in Stage II and 70% in Stage III and IV disease samples (Supplementary Figure 6D).

Seeking confirmation for the above finding, we prepared TMAs of a selection of tumor specimens from a non-consecutive cohort of 46 NMIBC and 42 MIBC FFPE patient samples representative of all tumor grade and stages (Supplementary Table 1) and evaluated protein expression of both CD44 and CD44v6. The resulting data revealed that 78.8% (88.1% of NMIBC and 68.4% from MIBC) of the assayed sections were positive for CD44v6 expression, restricted to the tumor area (Supplementary Figure 7A). Associations of CD44s and CD44v6 receptor expression with clinical data were explored (Figure 1B & 1C; Supplementary Figure 7B-7D). Even though significantly higher CD44s expression levels were detected among patients from advanced stage disease, no association between CD44v6 receptor expression and disease stage was found (Figure 1C; Supplementary Figure 7B). Nevertheless, a tendency towards higher CD44v6 expression levels in high grade disease could be observed, whereas no evidence of differential CD44s receptor expression between low-grade and high-grade disease samples was detected (Figure 1C). Additionally, samples from NMIBC patients showing disease progression after recurrence as well as MIBC patients showing metastatic disease displayed higher CD44v6 expression levels (Figure 1C; Supplementary Figure 7D). Increased CD44s expression levels among the primary tumors of metastatic patients were also observed (Figure 1C).

**Figure 1.**
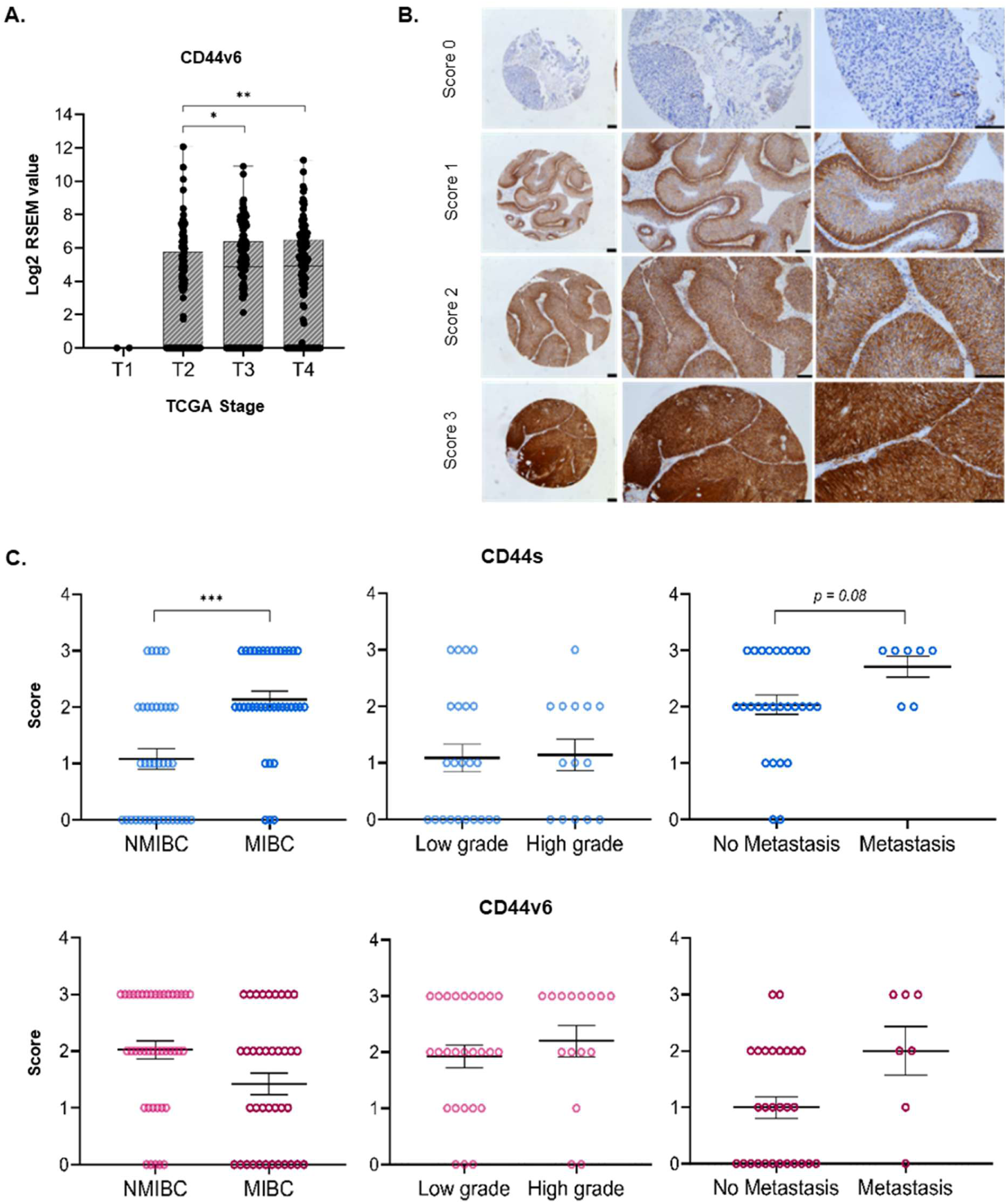
CD44 and CD44v6 expression analysis using BC patient samples. A) Graph showing relative *CD44v6* gene expression levels across 406 BC patients. Patients from stage I to stage IV disease were included in this analysis. The gene expression data were obtained from the TCGA database. Boxplots were used for representation, showing all individual values. B) Representative image of CD44v6 receptor staining among human NMIBC samples. CD44 and CD44v6 expression was scored (score 0 to 3) based on quantity and intensity of IHC staining (score 0: negative samples; score 3: complete and very intense staining). Scale bars, 100 μm. C) Evaluation of CD44s and CD44v6 receptor expression in BC patient samples based on clinical data, such as tumor invasiveness (NMIBC vs MIBC), grade (Low vs High) and the presence of metastasis. Error bars represent the mean ± Standard Error of the Mean (SEM). *p-value<0.05, **p-value<0.01, ***p-value<0.001.

### 3.2 Transcriptomic profiling based on CD44 and CD44v6 expression maps to specific molecular subtypes

FFPE samples from the cohort described above were utilized to characterize the transcriptomic landscape associated with CD44s and CD44v6-positive tumors. We selected 6 CD44s High (CD44s+; IHC score 3) and 6 CD44s negative (CD44s-; IHC score 0) tumors, as well as 8 CD44v6 High (CD44v6+; IHC score 3) and 3 CD44v6 negative (CD44v6-; IHC score 0) tumors for RNA sequencing (Figure 2). Analysis of the resulting RNAseq data by Principal Component Analysis (PCA) revealed a clear discrimination between CD44s- and CD44s+ samples as well as between CD44v6- and CD44v6+ samples (Figure 2A & 2B). Interestingly, the two CD44s-CD44v6-samples and the four CD44s-CD44v6+ samples were also clearly separated in CD44s-tumors (Figure 2A); a similar phenomenon was observed in CD44v6+ tumors, where CD44v6+CD44s- and CD44v6+CD44s+ samples were also easily discriminated (Figure 2B). These results were confirmed by unsupervised clustering analysis of differently expressed (DE) genes (Figure 2A’ & 2B’). Specifically, transcriptional analysis of CD44s+ and CD44s-tumors found 3061 DE gene transcripts, of which 1816 were up-regulated and 1245 were down-regulated in CD44s+ tumors compared to their CD44s-counterparts (Supplementary Data 2). Likewise, in CD44v6+ vs CD44v6-tumors there were 1295 DE gene transcripts, of which 815 were up-regulated and 480 were down-regulated in CD44v6+ samples compared to their CD44v6-counterparts (Supplementary Data 3). For CD44v6 specifically, the three distinct groups found by PCA were also found by unsupervised filtered clustering (Figure 2B’). Additionally, GSEA, GSVA and Enrichr analysis showed that CD44s+ tumors exhibited an enrichment in several biological pathways involved in tumor development and progression such as EMT, extracellular matrix (ECM) organization, inflammatory response, angiogenesis and proliferation (Supplementary Figure 8-10). CD44v6+ tumors showed enrichment in hypoxia, EMT and MYC signaling pathways as well as the activation of p53 and especially p63 signaling by GSEA and Enrichr analysis (Supplementary Figure 8 & 10). Interestingly, these cells also showed a decrease in the expression of genes downregulated in the absence of *CDH1* (E-cadherin) in tumor cells (Supplementary Figure 10).

We next set out to find if there was a correspondence between the groups defined from these RNAseq molecular profiling data and four different, previously existing molecular classification systems devised for BC. Indeed, we observed that the samples stratified by CD44s-/CD44s+ or CD44v6-/CD44v6+ transcriptional status did fit into groups previously defined by BC molecular classifiers (Figure 2C-2C’’). For instance, if the samples were separated according to CD44s expression, the resulting groups corresponded perfectly to the basal and luminal subtype groups defined by the UNC subtype classification (Kamoun et al., 2020) (Figure 2C). In turn, if the data were stratified according to expression of the CD44v6 isoform, tumors that were positive for CD44v6 but negative for CD44s mapped onto the mesenchymal-like/UroA subtypes of the LUND classifier (Kamoun et al., 2020) and the stromal rich/LumU tumor subtype of the Consensus MIBC classification (Kamoun et al., 2020) (Figure 2C’-2Ć’), without spilling over to other groups. Interestingly, CD44v6-tumors corresponded exactly to the GU tumor subtype of the LUND classifier and the LumP subtype of the Consensus MIBC classification (Figure 2C’-2Ć’). These findings were confirmed using the s265 cohort, showing that the v6 isoform constitutes a significant part of CD44 expression in BLCA and exhibits subtype-specificity distinct from the standard isoform CD44s. CD44v6 correlates more strongly with basal gene expression than other isoforms and its expression is more variable in higher tumor stages (Supplementary Figure 9).

**Figure 2.**
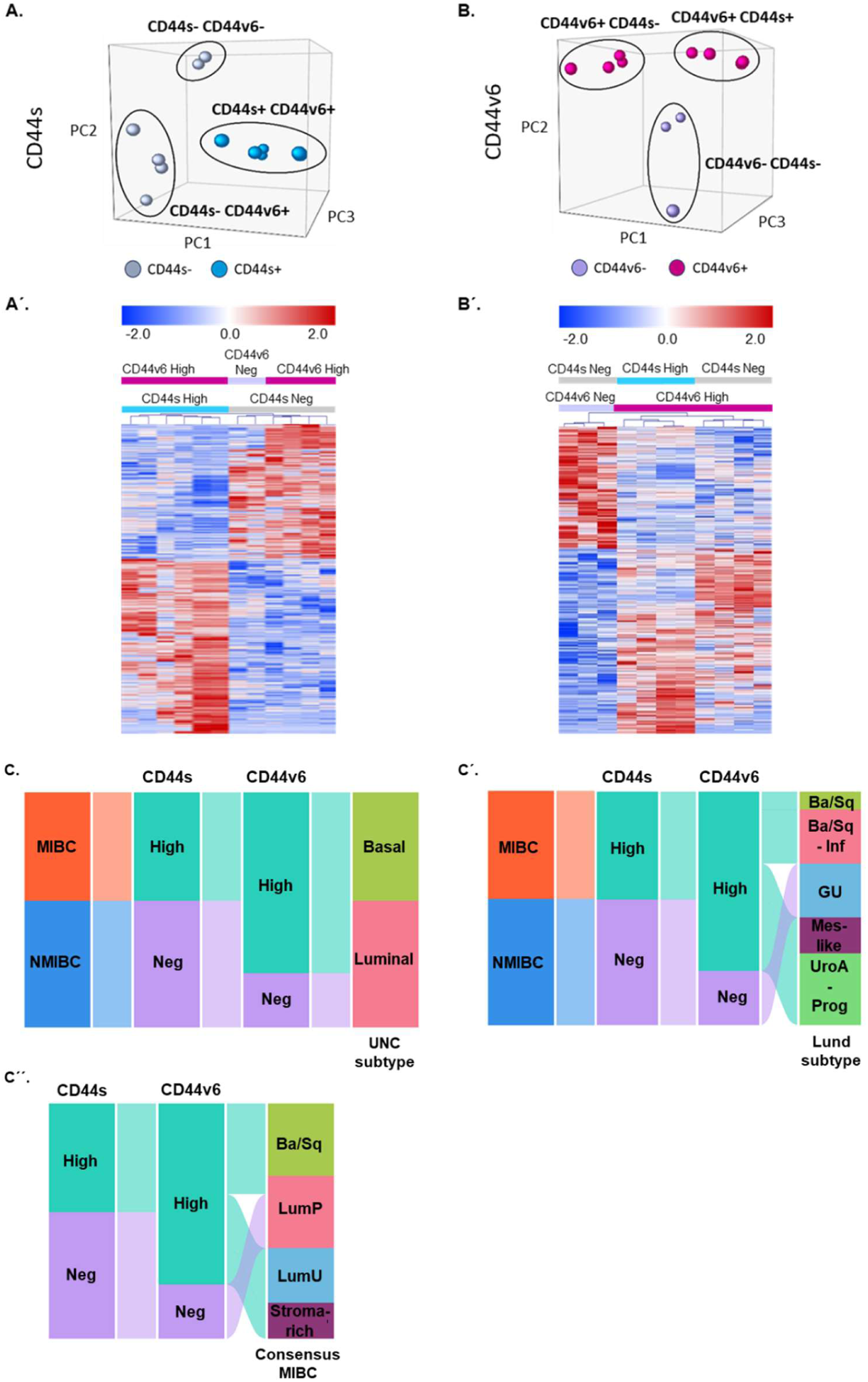
RNA-Seq analysis of selected BC patient samples based on CD44s and CD44v6 expression. A-Á) RNA-Seq outcomes comparing CD44s High (CD44s+) and CD44s negative (CD44s-) tumors, represented by PCA (A) and unsupervised clustering of DE genes visualized by a heatmap (Á). B-B’) RNA-Seq outcomes comparing CD44v6 High (CD44v6+) and CD44v6 negative (CD44v6-) tumors, represented by PCA (B) and unsupervised clustering of DE genes visualized by a heatmap (B’). C-Ć’) Alluvial plots showing CD44s- and CD44v6-expression based on sample distribution according to reported molecular BC classification systems (Kamoun et al., 2020): UNC classification (C), Lund classification (Ć), and Consensus MIBC classifier (Ć’). Ba/Sq = Basal/Squamous, Inf = Infiltrated, GU = Genomically Unstable, Mes-like = Mesenchymal-like, UroA-Prog = Urobasal A-Progressed, LumP = Luminal Papillary, LumU = Luminal Unstable.

### 3.3 CD44 and CD44v6 receptor density among human BC cell lines

We analyzed by flow cytometry ten human BC cell lines of known genomic characteristics representing different disease stages and BC subtypes (Supplementary Table 3) in order to examine whether their *in vitro* behavior was influenced by CD44 and/or CD44v6 surface expression status. As shown in Figure 3A, these lines exhibited highly variable CD44 receptor expression levels. A positive association was observed between CD44 receptor density and lines from more advanced disease stage or higher grade. CD44v6 receptor expression was found in all evaluated cell lines, although in contrast with the tumor transcriptomic data, elevated CD44v6 receptor expression levels were also observed among cell lines from lower stage and grade disease (Figure 3A). This phenomenon was not caused by differences in posttranslational processing, since *CD44v6* mRNA expression levels in these samples, as assayed by qPCR, agreed with the observed CD44v6 receptor expression (Supplementary Figure 11A). Significant differences between CD44 and CD44v6 receptor density were found for all BC cell lines, with cell lines representative of higher stage and grade tumors showing higher CD44:CD44v6 expression ratios (Figure 3A; Supplementary Figure 11B). Percentages of cells expressing CD44 and CD44v6 receptors were determined for all the abovementioned cell lines (Figure 3B; Supplementary Figure 12).

### 3.4 Generation and transcriptomic analysis of CD44 Low, CD44 High and CD44v6 positive cell lines

Based on their significantly different CD44 and CD44v6 receptor expression levels, the RT-112 and J82 cell lines were selected for further experimentation. Not only are CD44 and CD44v6 expression levels different between them, but the percentage of CD44v6-presenting cells is also markedly dissimilar (whereas approximately 97% of RT-112 cells and 100% of J82 cells are CD44+, 95% and 8% are CD44v6+, respectively) (Figure 3B).

CD44 Low, CD44 High and CD44v6 positive cell lines were successfully established from RT-112 and J82-derived FACS-isolated cell populations (Figure 3C & 3D; Supplementary Figure 1). Maintenance of the corresponding CD44 and CD44v6 positive phenotypes was verified by continuous subculturing and comparison of cultures at initial time or after 5 weeks of subculturing. The parental and newly established cell lines used in this study were named as following: RT-112 Parental, RT-112 CD44 Low, RT-112 CD44 High, RT-112 CD44v6 and J82 Parental, J82 CD44 Low, J82 CD44 High, J82 CD44v6 (Figure 3D).

We next compared RNAseq profiles between the generated cell lines (CD44 Low, CD44 High and CD44v6 from either RT-112 or J82) and their parental lines (RT-112 and J82) (Supplementary Data 4). The observed differences in *CD44* gene expression levels were concordant with the previously observed differences in CD44 protein expression (Figure 3E). Unsupervised classification, based on gene expression, clustered separately the parental and CD44 Low cell lines, whereas the CD44 High and CD44v6 lines clustered together (Supplementary Figure 13). As expected, basal gene expression appears elevated in CD44-high RT112 cells (Supplementary Data 4), indicating upregulated Cyclin D1/FOS/JUN pathway, basal lamina adhesion molecules (LAMA/B, ITGs), and NT5E, which are co-expressed with CD44. This suggests that variability in basal status is preserved in these cell lines and correlates positively with CD44 expression.

Analysis of upregulated genes shared between the CD44 High cell lines and CD44 High tumors detected a highly significant enrichment of genes involved in a variety of biological processes, including EMT, ECM interactions and cell migration (Figure 3F). When the same analysis was performed on upregulated genes shared between CD44v6 positive cell lines and CD44v6 High tumors, enrichment was again detected for genes of the p53 family (p53 and TAp63), which are known regulators of cell function that control various processes involved in cancer progression, including cell division, proliferation, genomic stability, cell cycle arrest, senescence, and apoptosis. Other enriched genes shared between CD44v6 cell lines and CD44v6 High tumors belonged to pathways involved in cell-cell and cell-ECM interactions (Figure 3F).

**Figure 3.**
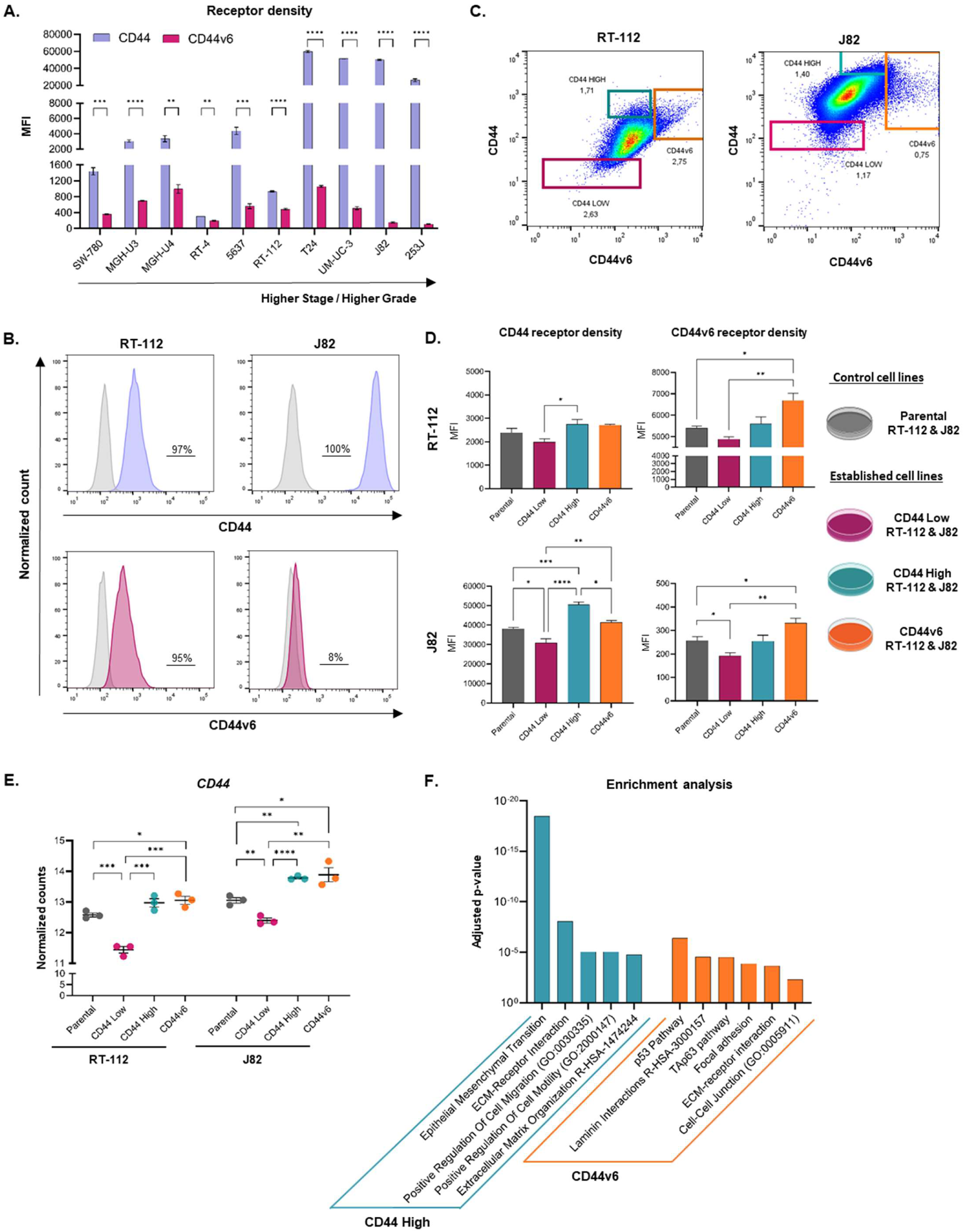
Generation and genomic analysis of cell lines with differential CD44 and CD44v6 expression. A) CD44 and CD44v6 receptor density among a broad variety of BC cell lines. B) Percentage of RT-112 and J82 cells expressing CD44 and CD44v6 receptors. C) Representative figures of FACS-isolated cell populations (CD44 Low, CD44 High and CD44v6 High) from RT-112 and J82 cell lines. D) CD44 and CD44v6 receptor density of newly established CD44 Low, CD44 High and CD44v6 cell lines from RT-112 and J82-derived FACS isolated cell populations. E) *CD44* gene expression levels across the newly generated cell lines, analyzed by RNA sequencing. Enrichment analysis on commonly upregulated genes across either CD44 High cell lines and CD44 High tumors or CD44v6 cell lines and CD44v6 High tumors, using Enrichr. The Median Fluorescence Intensity (MFI) allows for quantification of receptor expression. Error bars represent the mean ± SEM. *p-value<0.05, **p-value<0.01, ***p-value<0.001, ****p-value<0.0001.

### 3.5 Association of CD44 and CD44v6 expression with proliferation, migration and invasiveness

In order to examine whether the transcriptomic profiles of the CD44 Low, CD44 High and CD44v6 lines are recapitulated by their *in vitro* characteristics, we evaluated their proliferative, migrating and invading capabilities. Significant differences in proliferative capacity were observed between the parental, CD44 Low, CD44 High and CD44v6 cell lines *in vitro* (Figure 4A; Supplementary Figure 2B). RT-112 CD44 Low and J82 CD44 Low cell lines exhibited higher proliferative capacities than their respective parental, CD44 High and CD44v6 cell lines. Also, CD44 High cell lines were found to proliferate significantly slower than their respective parental and CD44 Low cell lines. Interestingly, the RT-112 CD44v6 cell line showed a significantly lower proliferative capacity than the RT-112 CD44 Low cell line but was found to proliferate considerably faster than the RT-112 CD44 High cell line. The J82 CD44v6 cell line behaved similarly, the only difference being that it exhibited no differences regarding cell proliferation with the J82 CD44 High cell line after 72 h of cell growth.

Evaluation of the migration and invasion capacity of the established cell lines also demonstrated differences between parental, CD44 Low, CD44 High and CD44v6 cell lines *in vitro* (Figure 4B-4B’ & 4C, Supplementary Figure 3 & 4). On the one hand, CD44 High cell lines, whether derived from the RT-112 or J82 parental cell line, exhibited a reduced ability to migrate when compared to their respective CD44v6 and parental cell lines. On the other hand, CD44 Low cell migration seemed to depend on the identity of the parental line. The migration capacity of RT-112 CD44 Low was similar to that of RT-112 CD44v6 but higher than that of RT-112 CD44 High and their parental cell line (Figure 4B-4B’; Supplementary Figure 3C). However, the migration capacity of J82 CD44 Low tended to be lower than that of J82 CD44v6 and their parental cell line. No significant difference was found between the migration capacity of the J82 CD44 Low and J82 CD44 High cell lines (Figure 4B’; Supplementary Figure 3C).

As for invasion capacity, the RT-112 CD44v6 cell line was significantly more invasive than its parental, CD44 Low and CD44 High cell lines; among them, RT-112 CD44 High was the least invasive. In contrast, the opposite was true for J82-derived cell lines, among which CD44 High exhibited a capacity for invasion significantly higher than that of the J82 CD44v6 and J82 CD44 Low cell lines (Figure 4C).

Interestingly, differences were found when comparing the CD44 and CD44v6 receptor phenotypes of the cell populations seeded in the Boyden chambers with those of the cells that had invaded the lower chamber by the end of the assay. In the J82 parental and its derived lines, the invading population exhibited a significantly higher proportion of CD44+/CD44v6+ cells than the original population (Figure 4D). Concordantly, the percentage of CD44+/CD44v6-cells was remarkably lower in the invading J82 cell populations (Supplementary Figure 4B). The same increase could not be observed in the RT-112-derived cell lines, as 90-97% of the starting population is already CD44v6+ (Supplementary Figure 4C).

**Figure 4.**
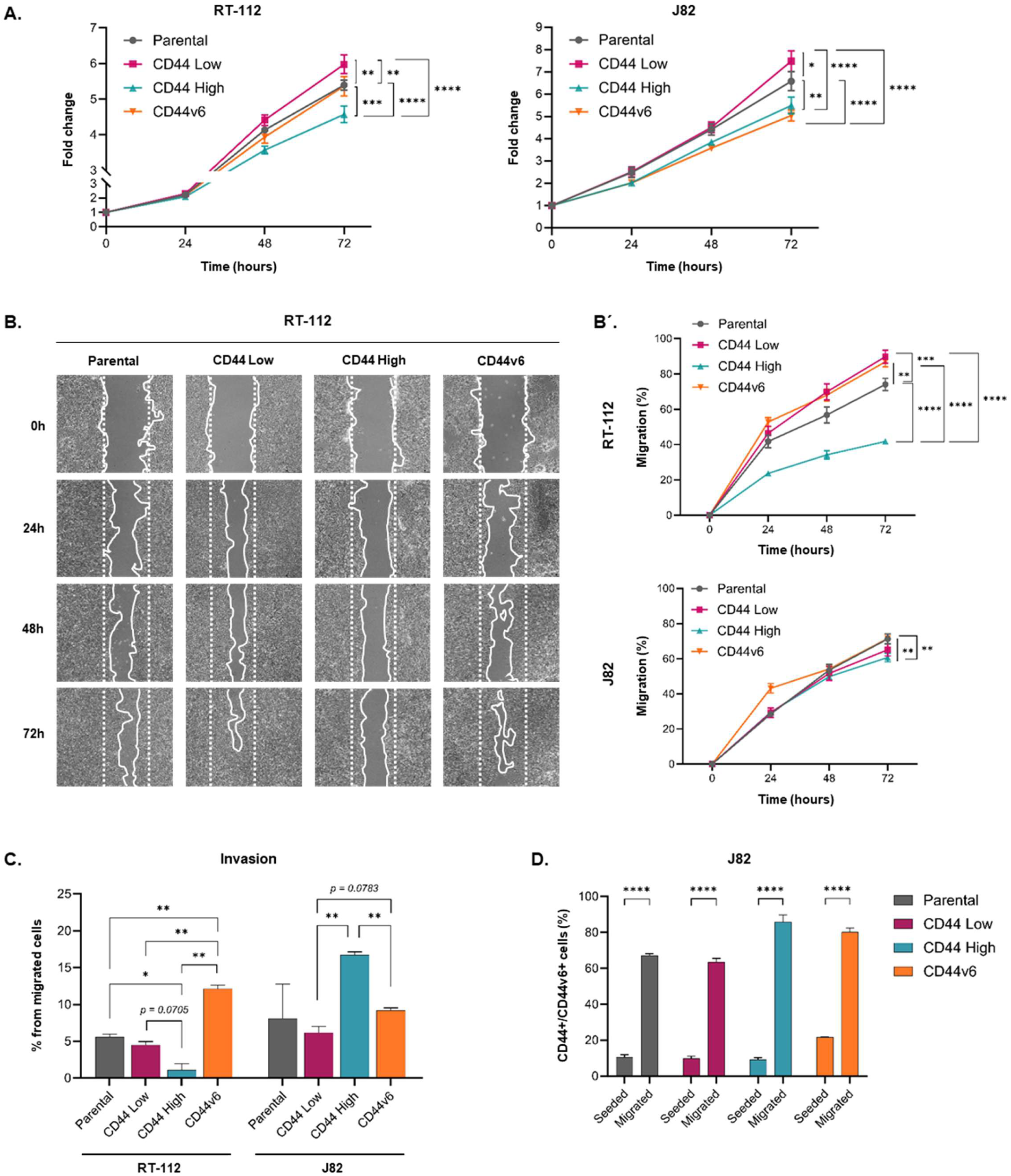
*In vitro* studies evaluating cell proliferation, migration, and invasion capacity of the CD44 Low, CD44 High and CD44v6 cell lines. A) Charts plotting the proliferation rate of parental (RT-112 and J82) and their respective CD44 Low, CD44 High and CD44v6 cell lines. Cell proliferation was measured 24 h, 48 h and 72 hours after seeding. The chart plots the fold change compared to time point 0. B-B’) Evaluation of the migration capacity of parental (RT-112 and J82) and their respective CD44 Low, CD44 High and CD44v6 cell lines. Representative image of RT-112 wound healing after 24 h, 48 h and 72 h of wound scratching (B). Charts plotting migration capacity, as represented by wound healing percentage 24 h, 48 h and 72 h after scratching (B’). C) Evaluation of the invasion capacity of parental (RT-112 and J82) and their respective CD44 Low, CD44 High and CD44v6 cell lines. Only cells with migration capacity were included in this analysis. The invasion capacity of each cell line is represented by the percentage of cells with invasive features. D) Percentage of CD44+/CD44v6+ cells in the seeded and invading J82 cell populations. Error bars represent the mean ± SEM. *p-value<0.05, **p-value<0.01, ***p-value<0.001, ****p-value<0.0001.

### 3.6 Increased in vivo tumorigenesis of established CD44v6 cell lines

The effect of CD44v6 on *in vivo* tumorigenesis was evaluated only with RT-112-derived cell lines, since the J82 parental line does not generate tumors in *nude* mice. RT-112 parental, as well as RT-112 CD44 Low, RT-112 CD44 High and RT-112 CD44v6 were subcutaneously injected into the flanks of immunodeficient *nude* mice, after which tumor implantation and growth were followed (Figure 5A).

With regards to tumor implantation, tumor development was observed in 100% (8/8) of animals receiving CD44 High xenografts, 88% (7/8) of animals receiving CD44v6 xenografts, and 63% (5/8) of the animals receiving the parental RT-112 cell line. Tumor implantation rate with CD44 Low xenografts was a very low 25% (2/8). Clear differences in tumor growth were observed, with the CD44v6 cell line being responsible for the fastest growing tumors (Figure 5B, Supplementary Figure 14A & 14B), followed by the parental and CD44 High cell lines, which exhibited an intermediate tumor growth rate, and trailed by CD44 Low with the slowest median rate of tumor growth (Figure 5B & 5C). These differences were confirmed upon tumor weighing after euthanasia (Supplementary Figure 14C).

Histological analysis uncovered major phenotypic differences between tumors derived from different cell lines (Figure 5D). Parental and CD44 Low-derived xenografts produced well differentiated tumors with a urothelial-like structure and a well-defined basal lamina, separating tumor and stroma. In contrast, CD44 High- and CD44v6-derived xenografts produced considerably less differentiated tumors of a more aggressive phenotype with no clear separation of tumor and stroma and cell infiltration into the stromal compartment, probably owing to the upregulation of genes involved in ECM interaction and the downregulation of genes associated to *CDH1* observed during transcriptomic analysis. When tumor cells were analyzed by flow cytometry, it was found that CD44v6-derived tumors exhibited significantly higher CD44 and CD44v6 receptor expression levels than the other cell lines (Figure 5E; Supplementary Figure 15).

Taking into account that transcriptomic analysis pointed at EMT as one of the processes associated with CD44 and CD44v6 expression, with a decrease in *CDH1* related genes in the case of CD44v6+ tumors and considering the more undifferentiated appearance of CD44 High and CD44v6-derived tumors, we decided to evaluate expression of *CDH1* in the tumor sections. We found a significantly decreased expression of *CDH1* among CD44v6-derived tumors, compared to parental-, CD44 Low- and CD44 High-derived tumors, further emphasizing the potential role of CD44v6 in tumor aggressiveness (Figure 5F & 5G).

**Figure 5.**
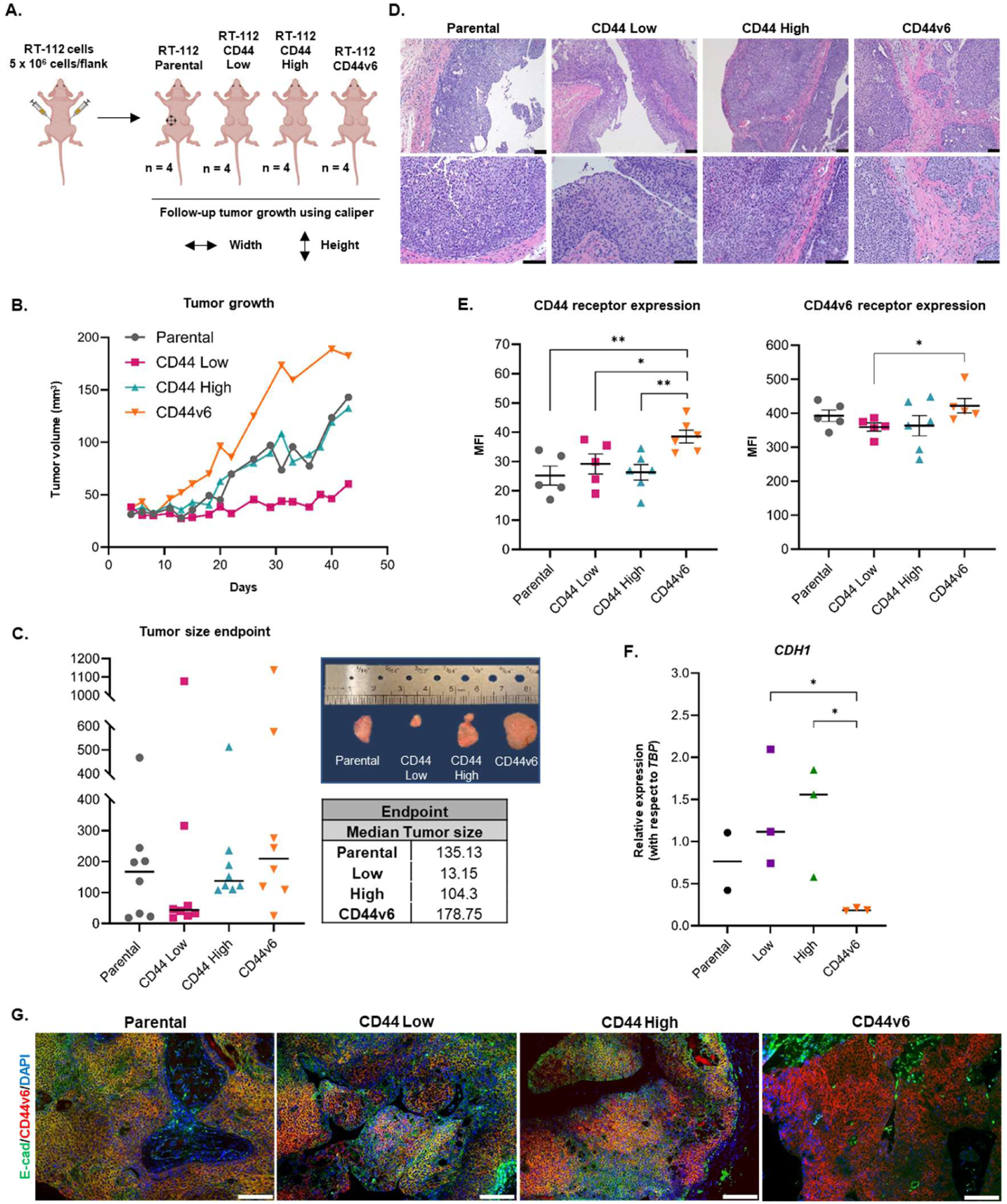
*In vivo* study evaluation of the tumorigenesis of CD44 Low, CD44 High and CD44v6 cell lines. A) Schematic representation of study design. The “n” represents the number of animals used in this study. B) Chart plotting the evolution of median tumor volume (mm^3^) with time. C) Individual tumor sizes at study endpoint. Tumor size is represented as tumor volume (mm^3^). Tumor sizes at the study endpoint were calculated by subtracting starting tumor size from final tumor size. Median tumor sizes at the study endpoint are also shown. D) Representative histological images of the tumors developed from RT-112 parental, CD44 Low, CD44 High and CD44v6 cell lines. Scale bars, 100 μm. E) CD44 and CD44v6 receptor expression in the developed tumors, analyzed by flow cytometry and the OMIQ Data Science Platform. Error bars represent the mean ± SEM. F) *CDH1* expression levels among the developed tumors, relative to the *TBP* housekeeping gene. Medians are also shown. Representative images of immunofluorescence (IF) analysis, showing E-cadherin expression (green) and CD44v6 expression (red) in the developed tumors. Nuclei are stained with DAPI (blue). Scale bars, 100 μm. The Median Fluorescence Intensity (MFI) allows for quantification of receptor expression. *p-value<0.05, **p-value<0.01.

### 3.7 Cisplatin sensitivity is decreased in established CD44v6 cell lines

In order to assess whether miRNA regulation was involved in a differential splicing switch towards CD44v6 expression, we compared the levels of 827 miRNAs in the parental and CD44v6 cell populations using nCounter miRNA analysis. Comparative analysis of these cell lines revealed a significant differential expression of 41 miRNAs, of which 14 miRNAs were upregulated and 27 miRNAs were downregulated in the parental cell lines compared to the CD44v6 cell lines (Supplementary Data 5). Unsupervised hierarchical clustering of these DE miRNAs significantly separated the parental cell lines from the CD44v6 sorted cell populations (Supplementary Figure 16A). Interestingly, by using the ROSALIND analysis system (NanoString Technologies), we found that these DE miRNAs are associated with the response to chemotherapeutic drugs such as cisplatin and doxorubicin (Supplementary Figure 16B).

Given the relevance of cisplatin-based therapy in the management of MIBC, we analyzed this aspect in depth. We used the cisplatin resistance database published by Huang et al. (2021) on the RNAseq data from CD44v6-expressing tumors and cell lines. In all cases, tumors and cell lines with high expression of CD44v6 exhibited a clear upregulation of genes from pathways related to cisplatin resistance such as CSCs, ECM interaction and EMT (Figure 3F; Supplementary Figure 16C).

To further support the role of CD44v6 in platinum sensitivity, we analyzed the expression of the CD44v6 receptor among cisplatin-remnant cells, both *in vitro* and *in vivo*. After *in vitro* treatment of RT-112 cells with cisplatin at the IC_50_ concentration, remnant tumor cells showed CD44v6 receptor expression (Figure 6A). Moreover, we used a previously generated RT-112 xenograft model ^38^ to monitor CD44v6 expression among cisplatin-remnant cells *in vivo*. In this experiment, animals were treated with a dose of 6 mg/kg cisplatin once a week when a tumor volume of 150-250 mm^3^ was reached ^38^. As shown in Figure 6B, remnant tumor cells after cisplatin treatment were indeed highly positive for CD44v6 expression.

With these results we decided to evaluate cisplatin sensitivity among the established cell lines. Moderate albeit significant differences in cisplatin sensitivity were found between the parental, CD44 Low, CD44 High and CD44v6 cell lines *in vitro* (Figure 6C). Interestingly, the CD44v6 cell lines showed significantly higher resistance to cisplatin treatment compared to parental and CD44 Low cell lines, independent of the original cell line. Additionally, the J82 CD44v6 line was significantly more resistant to cisplatin than J82 CD44 High. In general, J82-derived cell lines showed lower sensitivity to cisplatin than RT-112-derived cell lines, as reflected in their higher IC_50_ values.

**Figure 6.**
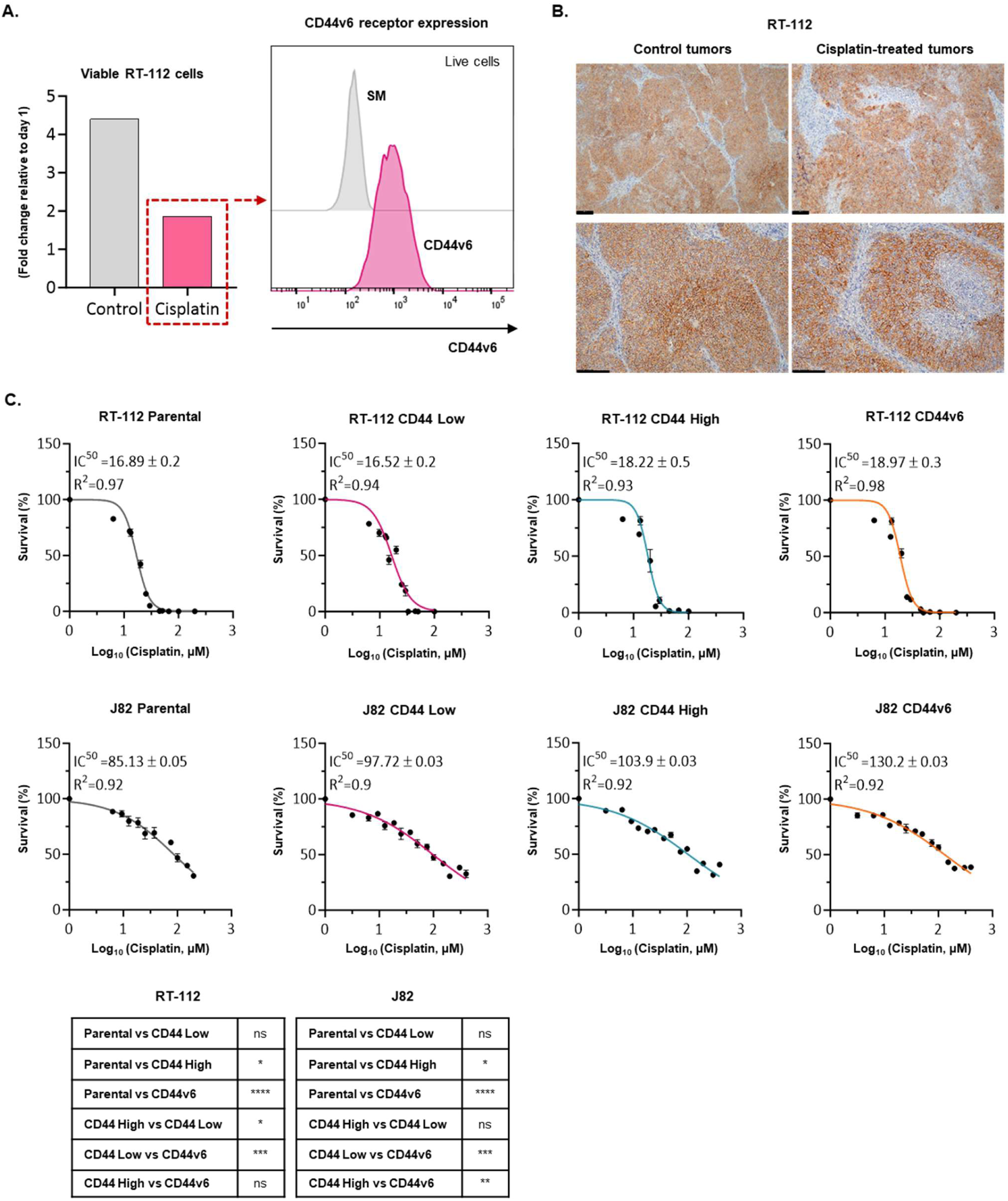
Cisplatin sensitivity of parental (RT-112 and J82) cell lines and their respective CD44 Low, CD44 High and CD44v6 cell lines. A) CD44v6 receptor expression in remnant RT-112 tumor cells following cisplatin treatment at the IC50 concentration. B) Representative IHC images of CD44v6 expression among non-treated (control) and cisplatin-treated tumors. Scale bars, 100 μm. C) *In vitro* study of cell viability by XTT. Cell survival is represented as percentages. Cisplatin concentrations are in Log10 scale. Tables showing the level of statistical significance of differences in cisplatin sensitivity between the parental cell lines (RT-112 and J82) and their respective CD44 Low, CD44 High and CD44v6 cell lines. ns = not significant, *p-value<0.05, **p-value<0.01, ***p-value<0.001, ****p-value<0.0001.

## 4. DISCUSSION

CD44 is a hyaluronan-binding cell surface adhesion molecule involved in tumor development and metastasis that supports the formation of a favorable tumor microenvironment and the expression of a chemotherapy-resistant phenotype ^39,40^. Its overexpression has been shown to attenuate the cytotoxicity of chemotherapy in various cancers, resulting in poor patient prognosis ^40^. Important as its role in tumorigenesis might be, the tissue distribution of CD44 is far too wide for this molecule to be used in targeted cancer therapies, as the latter relies on the exclusive or augmented expression of the target on tumoral cells.

CD44 transcripts may undergo alternative splicing of the exons coding for its extracellular domain, generating several highly glycosylated CD44 variants (CD44v) that adopt new functionalities and exhibit a much more specific tissue distribution than the standard isoform ^41–47^. One such variant is CD44v6, the isoform containing the variable exon 11-encoding region, whose expression is restricted to subsets of epithelia and specific developmental stages ^48^. CD44v6 is overexpressed in several cancers, including HNSCC, ovarian cancer, colorectal cancer, and BC ^23,49–51^. High CD44v6 expression is often associated with poor prognosis, aggressive disease, and increased invasive and metastatic potential ^52–54^.

Due to its limited expression in healthy tissues and its marked overexpression in various cancers, CD44v6 has been considered a promising therapeutic target for cancer, even reaching clinical trials as a target for molecular radiotherapy or CAR-T cells in HNSCC ^55,56^. In this study we focused on the biology of CD44 and its variant CD44v6 in BC, as part of an effort to understand its role in the pathophysiology of this disease and to provide data informing its appraisal as a potential prognostic marker and therapeutic target in this clinical setting.

We first examined CD44v6 expression in BC samples from different stages by using TCGA transcriptomic data and FFPE patient samples studied by IHC. Remarkably, we found CD44v6 expression in more than 75% of all tumors independently of grade and stage, which was particularly high in metastatic forms of the disease. These findings coincide with those of Z. Wang et al. (2018), who inferred that CD44s/CD44v6 play a fundamental role not only on the generation of cancer initiation cells but also in subsequent steps of the metastatic cascade.

Our findings suggest that CD44v6 facilitates tumor progression through enhanced cell migration, invasion, and interaction with the tumor microenvironment, which is consistent with previous research on the role of CD44 variants in cancer metastasis ^58^. The same has previously been reported for ovarian ^59^, colorectal ^60^ and breast cancer ^61^, where several research works and meta-analyses have validated CD44v6 as a reliable marker of poor prognosis. The specific role of the v6 region was first demonstrated by Günthert et al. (1991), who identified this epitope and demonstrated that overexpression of the v6 isoform in non-metastasizing BSp73AS cells suffices to establish full metastatic behavior. After that, many individual studies have reported that the v6 isoform interacts with integral membrane and cytosolic signaling molecules as well as proteases and transcriptional regulators involved in migration and invasion. CD44v6 promotes MET activation, which requires the interaction of the CD44 cytoplasmic tail with ERM proteins for activation of the Ras-MAPK pathway ^63^. In ovarian cancer, the CD44v6-ECM interaction also promotes PI3K pathway activation, MET transcription ^64^ and metastasis by mediating cell invasion into the peritoneum ^18^ in a process suggested to induce NF-κB activation. This characteristic may serve an important role in the potential use of chemotherapeutic drugs that target this critical molecular mechanism for the treatment of cancer.

The transcriptomic analyses presented here of patient-derived tumors as well as tumor-derived cell lines revealed that CD44 and CD44v6 are involved in the EMT process and the interaction of tumoral cells with the extracellular matrix during BC. CD44 has been previously shown to play these roles in other tumor types ^18,58^. In the particular case of CD44v6, previous research has demonstrated that the secretion of hepatocyte growth factor (HGF), osteopontin (OPN), and stromal-derived factor 1α (SDF-1) by tumor-associated cells increases CD44v6 expression in colorectal CSCs via activation of the Wnt/β-catenin pathway, which promotes migration and metastasis ^58^. These authors reported that CD44v6 negative progenitor cells do not give rise to metastatic lesions but, when treated with cytokines, acquire CD44v6 expression and metastatic capacity. Importantly, phosphatidylinositol 3-kinase inhibition selectively killed CD44v6 colorectal CSCs and reduced metastatic growth. This will be very relevant for the design of future therapeutic strategies targeting CD44v6-expressing cells with specific PI3K signaling inhibitors.

Regarding the molecular classification of tumors expressing CD44v6, we found that unsupervised clustering of DE genes from transcriptomics data separated BC tumors into groups according to their CD44v6 status and, importantly, that these groups fit very well onto the LUND and Consensus classification schemes. This finding has important clinical implications, as the molecular subtypes of BC tumors correlate well with pathological response and overall survival. The CD44v6 positive tumors grouped together by unsupervised clustering corresponded to the Basal/Squamous subgroup, which has been shown to exhibit the worst outcome in three different cohorts ^65^. In this regard, Sjödahl et al. (2022) found that tumors with high expression of *SPP1* (osteopontin) were associated with poor response to chemotherapy, and osteopontin has been shown to increase the metastatic potential of hepatocellular cancer cells by raising plasma membrane CD44v6 expression and, therefore, cell adhesion to hyaluronic acid ^66^. Hence, the combination of SPP1 expression with a molecular classification could be used as a biomarker in future CD44v6-targeted therapies, since we demonstrated that high CD44v6-expressing cells exhibit increased resistance to cisplatin. Our findings are also in concordance with previous research on gastric cancer ^67^ demonstrating that CD44v6 expression increases the survival of gastric cancer cells in response to cisplatin, and that these cells override CD44v6-negative cells after cisplatin treatment. Similar results have been described in other cancer types ^68,69^. By interfering with the interaction of CD44v6 with its ligands or downstream signaling pathways, it may be possible to inhibit BC progression and overcome resistance to conventional treatments. However, therapeutic exploitation of CD44v6 requires a deeper understanding of its regulatory networks and the identification of specific inhibitors that can selectively target this variant without interfering with the normal physiological functions of CD44.

Limitations of our study include the retrospective nature of the patient cohort and the reliance on immunohistochemical analysis for protein expression, which may not fully capture the dynamic and complex regulation of CD44 and its variants. Future research should aim to validate these findings in prospective studies and explore the therapeutic potential of targeting CD44v6 using *in vivo* models. In addition, investigating the role of CD44v6 in the tumor microenvironment and its interaction with immune cells may provide further insights into its function in tumor immunity and suggest novel combinatorial therapeutic strategies.

## 5. CONCLUSIONS

In conclusion, our study adds to the growing body of evidence supporting the role of CD44 and its variant CD44v6 in BC progression and highlights their potential as targets for therapeutic intervention. We showed that the expression of the CD44v6 isoform increases BC cell survival in response to cisplatin treatment. The development of CD44v6-targeted therapies could represent a significant advance in the treatment of BC, offering hope for improved patient outcomes.

## Supporting information

Manuscript with figures

## LIST OF ABBREVIATIONS

BC: Bladder cancer
NMIBC: non-muscle-invasive BC
MIBC: muscle-invasive BC
mBC: metastatic stage
ADC: antibody-drug conjugates
TAA: tumor-associated antigens
CSCs: cancer stem cells
HNSCC: head and neck squamous cell carcinoma
TCGA: Cancer Genome Atlas
RNA Seq: Ribonucleic Acid sequencing
BLCA: Bladder Urothelial Carcinoma
RSEM: RNA-Seq by Expectation-Maximization
RetBioH: Spanish Hospital Biobanks Network
TAM: tissue microarray
FFPE: formalin-fixed paraffin-embedded
HS: horse serum
FBS: fetal bovine serum
MFI: Median Fluorescence Intensity
FMO: Fluorescence Minus One
PCA: Principal component analysis
GSVA: Gene set variation analysis
EMT: Epithelial-Mesenchymal transition
ECM: Extracellular matrix
CD44v: CD44 variants
HGF: hepatocyte growth factor
OPN: osteopontin
SDF-1: stromal-derived factor 1α

## DECLARATIONS

### Ethics approval and consent to participate

Patient enrollment and data collection procedures were supervised and approved by the Ethical Committee for Clinical Research of the 12 de Octubre university hospital (CEIC16-011). Written informed consent was obtained from all patients/participants prior to the study.

All the animal experimental procedures were conducted according to the European and Spanish regulations in the field: European convention ETS 123, regarding the use and protection of vertebrate mammals used in experimentation and other scientific purposes, and Directive 2010/63/UE, Spanish Law 6/2013 and R.D. 53/2013 regarding the protection and use of animals in scientific research. All procedures were approved by the CIEMAT Animal Experimentation Ethics Committee in accordance with external and internal biosafety and bioethics guidelines, and were approved by the Consejería de Medio Ambiente, Agricultura e Interior de la Comunidad de Madrid (Spain; protocol numbers PROEX 150.8/21). Mice strains were housed at the CIEMAT laboratory Animal Facility (registration number ES280790000183).

### Consent for publication

All authors have approved the manuscript for submission, and we confirm that the content of the manuscript has not been published or submitted for publication elsewhere.

### Availability of data and materials

The datasets generated and analyzed during the current study are available in the GEO repository. GEO accession numbers are: GSE276685 and GSE276545.

### Authors’ Contributions

I.L. conducted all *in vivo* and cell-based experimental studies, contributed to the design of the study, analyzed and interpreted the data and drafted and revised the manuscript. C.R. contributed to the design of the study, analyzed and interpreted the cytomic and transcriptomic data and drafted and revised the manuscript. P.E. analyzed and interpreted the transcriptomic data on validation cohorts. I.A., L.G., A.MdB., M.A. and E.M. conducted cell-based experimental studies. L.M. and C.S. conducted *in vivo* experimental studies, interpreted the data and revised the manuscript. O.A. and R.S. contributed to the analysis of cytomic data. F.G., D.C., L.P., R.G. and J.L.R. contributed with the patient samples selection, tissue characterization, clinical data and TMA construction. G.S. contributed to the design of the study, analyzed and interpreted the data and revised the manuscript. J.M.P. and M.D. contributed to the design of the study, analyzed and interpreted the data and drafted and revised the manuscript.

## Acknowledgements

We thank the Histology Laboratory from CIEMAT, namely P. Hernandez Lorenzo, for the histological processing of tumor samples and A. M. Martín Dunn for his conscientious scientific and grammatical revision of the manuscript. J.M.P discloses support for the research of this work from European Regional Development Fund for Science and Innovation (SAF2015-66015-R and PID2019-110758RB-I00) and Instituto de Salud Carlos III (CIBERONC no. CB16/12/00228). M.D. discloses support for the research of this work from Instituto de Salud Carlos III (PI20/00813, DTS20/00043, DTS22/00002 and AC22/00015), and Scientific Foundation of the Spanish Association Against Cancer (FCAECC) (TRNSC213883DUEN and Transcan-3 JTC2022). C.R. discloses support for the research of this work from Fundación Eugenio Rodríguez Pascual (FERP-2022-79). I.L. is supported by a predoctoral fellowship from AECC (Spanish Ass. against Cancer), Predoctoral AECC 2019 grant number PRDMA19024LODE. L.M. is supported by Fundación Científica Asociación Española Contra el Cáncer (AECC), Postdoctoral AECC 2019 grant number POSTD19036MORA.

## SUPPLEMENTARY DATA

**Supplementary Data 1. Overview of TCGA data.** os = overall survival.

**Supplementary Data 2. Counts & DESEQ2 results based on CD44 differential expression across FFPE patient samples, obtained by RNAseq.**

**Supplementary Data 3. Counts & DESEQ2 results based on CD44v6 differential expression across FFPE patient samples, obtained by RNAseq.**

**Supplementary Data 4. Counts & DESEQ2 results comparing the generated cell lines, obtained by RNAseq.**

**Supplementary Data 5. Differentially expressed miRNAs between parental and generated CD44v6 cell lines.** Normalized values are obtained after nCounter miRNA analysis using the mean of housekeeping genes.

## REFERENCES

1. Sung, H. et al. Global Cancer Statistics 2020: GLOBOCAN Estimates of Incidence and Mortality Worldwide for 36 Cancers in 185 Countries. CA Cancer J Clin 71, 209–249 (2021).

2. Bray Bsc, F., et al. Global cancer statistics 2022: GLOBOCAN estimates of incidence and mortality worldwide for 36 cancers in 185 countries. (2024) doi:10.3322/caac.21834.

3. Sanli, O. et al. Bladder cancer. Nat Rev Dis Primers 3, (2017).

4. Lidagoster, S., Ben-David, R., De Leon, B. & Sfakianos, J. P. BCG and Alternative Therapies to BCG Therapy for Non-Muscle-Invasive Bladder Cancer. Curr Oncol 31, 1063–1078 (2024).

5. Von der Maase, H., et al. Gemcitabine and cisplatin versus methotrexate, vinblastine, doxorubicin, and cisplatin in advanced or metastatic bladder cancer: results of a large, randomized, multinational, multicenter, phase III study. J Clin Oncol 18, 3068–3077 (2000).

6. Patel, M. R. et al. Avelumab in metastatic urothelial carcinoma after platinum failure (JAVELIN Solid Tumor): pooled results from two expansion cohorts of an open-label, phase 1 trial. Lancet Oncol 19, 51–64 (2018).

7. Powles, T. et al. Efficacy and Safety of Durvalumab in Locally Advanced or Metastatic Urothelial Carcinoma: Updated Results From a Phase 1/2 Open-label Study. JAMA Oncol 3, (2017).

8. Rosenberg, J. E. et al. Atezolizumab in patients with locally advanced and metastatic urothelial carcinoma who have progressed following treatment with platinum-based chemotherapy: a single-arm, multicentre, phase 2 trial. Lancet 387, 1909–1920 (2016).

9. Sharma, P. et al. Nivolumab in metastatic urothelial carcinoma after platinum therapy (CheckMate 275): a multicentre, single-arm, phase 2 trial. Lancet Oncol 18, 312–322 (2017).

10. Bellmunt, J. et al. Pembrolizumab as Second-Line Therapy for Advanced Urothelial Carcinoma. N Engl J Med 376, 1015–1026 (2017).

11. Powles, T. et al. Enfortumab Vedotin and Pembrolizumab in Untreated Advanced Urothelial Cancer. N Engl J Med 390, 875–888 (2024).

12. Zschäbitz, S. et al. Enfortumab Vedotin in Metastatic Urothelial Carcinoma: Survival and Safety in a European Multicenter Real-world Patient Cohort. Eur Urol Open Sci 53, 31–37 (2023).

13. Aggen, D. H., Chu, C. E. & Rosenberg, J. E. Scratching the Surface: NECTIN-4 as a Surrogate for Enfortumab Vedotin Resistance. Clin Cancer Res 29, 1377–1380 (2023).

14. Chen, C., Zhao, S., Karnad, A. & Freeman, J. W. The biology and role of CD44 in cancer progression: therapeutic implications. Journal of Hematology & Oncology 2018 11:1 11, 1–23 (2018).

15. Xu, H. et al. The role of CD44 in epithelial–mesenchymal transition and cancer development. Onco Targets Ther 8, 3783 (2015).

16. Primeaux, M., Gowrikumar, S. & Dhawan, P. Role of CD44 Isoforms in Epithelial-Mesenchymal Plasticity and Metastasis. Clin Exp Metastasis 39, 391 (2022).

17. Mack, B. & Gires, O. CD44s and CD44v6 Expression in Head and Neck Epithelia. PLoS One 3, e3360 (2008).

18. Tjhay, F. et al. CD44 variant 6 is correlated with peritoneal dissemination and poor prognosis in patients with advanced epithelial ovarian cancer. Cancer Sci 106, 1421 (2015).

19. Kataki, A. et al. Membranous CD44v6 is upregulated as an early event in colorectal cancer: Downregulation is associated with circulating tumor cells and poor prognosis. Oncol Lett 22, (2021).

20. Hosseini, A. et al. The clinical significance of CD44v6 in malignant and benign primary bone tumors. BMC Musculoskelet Disord 24, 1–13 (2023).

21. Burman, D. R., Das, S., Das, C. & Bhattacharya, R. Alternative splicing modulates cancer aggressiveness: role in EMT/ metastasis and chemoresistance. 48, 897–914 (2021).

22. Mortensen, A. C. L. et al. Selection, characterization and in vivo evaluation of novel CD44v6-targeting antibodies for targeted molecular radiotherapy. Scientific Reports 2023 13:1 13, 1–11 (2023).

23. Omran, O. M. & Ata, H. S. CD44s and CD44v6 in diagnosis and prognosis of human bladder cancer. Ultrastruct Pathol 36, 145–152 (2012).

24. Sun, W. et al. TSVdb: A web-tool for TCGA splicing variants analysis. BMC Genomics 19, 1–7 (2018).

25. Cotillas, E. A. et al. A Versatile and Upgraded Version of the LundTax Classification Algorithm Applied to Independent Cohorts. J Mol Diagn 26, (2024).

26. Earl, J. et al. The UBC-40 Urothelial Bladder Cancer cell line index: a genomic resource for functional studies. BMC Genomics 16, (2015).

27. Love, M. I., Huber, W. & Anders, S. Moderated estimation of fold change and dispersion for RNA-seq data with DESeq2. Genome Biol 15, (2014).

28. R Core Team. R: A language and environment for statistical computing. R Foundation for Statistical Computing. (2022).

29. Saeed, A. I. et al. TM4: a free, open-source system for microarray data management and analysis. Biotechniques 34, 374–378 (2003).

30. Liberzon, A. et al. The Molecular Signatures Database (MSigDB) hallmark gene set collection. Cell Syst 1, 417 (2015).

31. Subramanian, A. et al. Gene set enrichment analysis: a knowledge-based approach for interpreting genome-wide expression profiles. Proc Natl Acad Sci U S A 102, 15545– 15550 (2005).

32. Hänzelmann, S., Castelo, R. & Guinney, J. GSVA: Gene set variation analysis for microarray and RNA-Seq data. BMC Bioinformatics 14, 1–15 (2013).

33. Kamoun, A. et al. A Consensus Molecular Classification of Muscle-invasive Bladder Cancer. Eur Urol 77, 420–433 (2020).

34. Tang, D. et al. SRplot: A free online platform for data visualization and graphing. PLoS One 18, (2023).

35. Schindelin, J., et al. Fiji: an open-source platform for biological-image analysis. Nat Methods 9, 676–682 (2012).

36. Kamoun, A. et al. A Consensus Molecular Classification of Muscle-invasive Bladder Cancer. Eur Urol 77, 420–433 (2020).

37. Huang, D. et al. A highly annotated database of genes associated with platinum resistance in cancer. Oncogene 2021 40:46 40, 6395–6405 (2021).

38. Rubio, C. et al. CDK4/6 Inhibitor as a Novel Therapeutic Approach for Advanced Bladder Cancer Independently of RB1 Status. Clin Cancer Res 25, 390–402 (2019).

39. Prochazka, L., Tesarik, R. & Turanek, J. Regulation of alternative splicing of CD44 in cancer. Cell Signal 26, 2234–2239 (2014).

40. Yaghobi, Z. et al. The role of CD44 in cancer chemoresistance: A concise review. Eur J Pharmacol 903, (2021).

41. Dixit, G., Zheng, Y., Parker, B. & Wen, J. Identification of Alternative Splicing and Polyadenylation in RNA-seq Data. J Vis Exp 2021, (2021).

42. El Marabti, E. & Younis, I. The Cancer Spliceome: Reprograming of Alternative Splicing in Cancer. Front Mol Biosci 5, (2018).

43. Li, Z. X. et al. Comprehensive characterization of the alternative splicing landscape in head and neck squamous cell carcinoma reveals novel events associated with tumorigenesis and the immune microenvironment. Theranostics 9, 7648–7665 (2019).

44. Oltean, S. & Bates, D. O. Hallmarks of alternative splicing in cancer. Oncogene 33, 5311– 5318 (2014).

45. Roy Burman, D., Das, S., Das, C. & Bhattacharya, R. Alternative splicing modulates cancer aggressiveness: role in EMT/metastasis and chemoresistance. Mol Biol Rep 48, 897–914 (2021).

46. Shen, S. et al. rMATS: robust and flexible detection of differential alternative splicing from replicate RNA-Seq data. Proc Natl Acad Sci U S A 111, E5593–E5601 (2014).

47. Wculek, S. K. et al. Dendritic cells in cancer immunology and immunotherapy. Nat Rev Immunol 20, 7–24 (2020).

48. Heider, K. H., Kuthan, H., Stehle, G. & Munzert, G. CD44v6: a target for antibody-based cancer therapy. Cancer Immunol Immunother 53, 567 (2004).

49. Bellerby, R. et al. Overexpression of Specific CD44 Isoforms Is Associated with Aggressive Cell Features in Acquired Endocrine Resistance. Front Oncol 6, (2016).

50. Orian-Rousseau, V. CD44, a therapeutic target for metastasising tumours. Eur J Cancer 46, 1271–1277 (2010).

51. Zhang, H. F., Hu, P. & Fang, S. Q. Understanding the role of CD44V6 in ovarian cancer. Oncol Lett 14, 1989–1992 (2017).

52. Escobar-Hoyos, L., Knorr, K. & Abdel-Wahab, O. Aberrant RNA Splicing in Cancer. Annu Rev Cancer Biol 3, 167 (2019).

53. Sun, Y., Yan, L., Guo, J., Shao, J. & Jia, R. Downregulation of SRSF3 by antisense oligonucleotides sensitizes oral squamous cell carcinoma and breast cancer cells to paclitaxel treatment. Cancer Chemother Pharmacol 84, 1133–1143 (2019).

54. Wang, B. D. & Lee, N. H. Aberrant RNA Splicing in Cancer and Drug Resistance. Cancers (Basel) 10, (2018).

55. Haist, C. et al. CD44v6-targeted CAR T-cells specifically eliminate CD44 isoform 6 expressing head/neck squamous cell carcinoma cells. Oral Oncol 116, 105259 (2021).

56. Spiegelberg, D. & Nilvebrant, J. CD44v6-Targeted Imaging of Head and Neck Squamous Cell Carcinoma: Antibody-Based Approaches. Contrast Media Mol Imaging 2017, (2017).

57. Wang, Z., Zhao, K., Hackert, T. & Zöller, M. CD44/CD44v6 a reliable companion in cancer-initiating cell maintenance and tumor progression. Front Cell Dev Biol 6, 397013 (2018).

58. Todaro, M. et al. CD44v6 is a marker of constitutive and reprogrammed cancer stem cells driving colon cancer metastasis. Cell Stem Cell 14, 342–356 (2014).

59. Wang, Y., Yang, X., Xian, S. H. U., Zhang, L. I. & Cheng, Y. CD44v6 may influence ovarian cancer cell invasion and migration by regulating the NF-κB pathway. Oncol Lett 18, 298 (2019).

60. Wang, J. L. et al. CD44v6 overexpression related to metastasis and poor prognosis of colorectal cancer: A meta-analysis. Oncotarget 8, 12866 (2017).

61. Hu, S. et al. CD44v6 targeted by miR-193b-5p in the coding region modulates the migration and invasion of breast cancer cells. J Cancer 11, 260–271 (2020).

62. Günthert, U. et al. A new variant of glycoprotein CD44 confers metastatic potential to rat carcinoma cells. Cell 65, 13–24 (1991).

63. Orian-Rousseau, V. et al. Hepatocyte Growth Factor-induced Ras Activation Requires ERM Proteins Linked to Both CD44v6 and F-Actin. Mol Biol Cell 18, 76 (2007).

64. Krause, D. S. & Van Etten, R. A. Tyrosine kinases as targets for cancer therapy. N Engl J Med 353, 172–187 (2005).

65. Sjödahl, G. et al. Different Responses to Neoadjuvant Chemotherapy in Urothelial Carcinoma Molecular Subtypes. Eur Urol 81, 523–532 (2022).

66. Gao, C. et al. Osteopontin-dependent CD44v6 expression and cell adhesion in HepG2 cells. Carcinogenesis 24, 1871–1878 (2003).

67. Pereira, C. et al. Expression of CD44v6-Containing Isoforms Influences Cisplatin Response in Gastric Cancer Cells. Cancers (Basel) 12, (2020).

68. Mooney, K. L. et al. The role of CD44 in glioblastoma multiforme. J Clin Neurosci 34, 1–5 (2016).

69. Lv, L. et al. Upregulation of CD44v6 contributes to acquired chemoresistance via the modulation of autophagy in colon cancer SW480 cells. Tumour Biol 37, 8811–8824 (2016).

