## Supplementary material for "CD44v6 Drives Tumor Aggressiveness and Chemoresistance in Bladder Cancer": Manuscript with figures

a. Cellular and Molecular Oncology and Genitourinary Tumor Group. Institute of Biomedical Research, Hospital Universitario 12 de Octubre, Madrid, Spain.
b. Molecular and Translational Oncology Division. Centro de Investigaciones Energéticas, Medioambientales y Tecnológicas (CIEMAT), Madrid, Spain.
c. Centro de Investigación Biomédica en Red de Cáncer (CIBERONC), Madrid, Spain.
d. Division of Oncology, Department of Clinical Sciences, Lund University, Lund, Sweden.
e. Division of Hematopoietic Innovative Therapies, Centro de Investigaciones Energéticas, Medioambientales y Tecnológicas (CIEMAT) and Centro de Investigación Biomédica en Red de Enfermedades Raras (CIBER-ER), Madrid, Spain.
f. Advanced Therapy Unit, Instituto de Investigación Sanitaria Fundación Jiménez Díaz (IIS-FJD/UAM), Madrid, Spain.
g. Uro-Oncology Unit, Department of Urology, Hospital Universitario 12 de Octubre, Madrid, Spain.
h. Pathology Department, Hospital Universitario 12 de Octubre, Madrid, Spain.
i. Department of Translational Medicine, Lund University, Malmö, Sweden.
j. Department of Urology, Skåne University Hospital, Malmö, Sweden.

### SUPPLEMENTARY INFORMATION

**Table of Contents**

**Supplementary Tables ........ 2**
**Supplementary Figures ........ 6**

#### Supplementary Tables

**Supplementary Table 1. Characteristics of patient cohorts.** PUNLMP = Papillary urothelial neoplasm of low malignant potential. Progression is defined as the appearance of MIBC in an initial NMIBC patient.

| Characteristics | No. of patients | % Patients |
|---|---|---|
| **Gender** | | |
| ***Male*** | | |
| **NMIBC** | 32 | 51% |
| **MIBC** | 31 | 49% |
| **Total** | 63 | |
| ***Female*** | | |
| **NMIBC** | 13 | 57% |
| **MIBC** | 10 | 43% |
| **Total** | 23 | |
| ***Unknown*** | | |
| **NMIBC** | 1 | 50% |
| **MIBC** | 1 | 50% |
| **Total** | 2 | |
| **Age (average)** | | |
| ***Male*** | 72.46 (range 51-93) | |
| ***Female*** | 69.33 (range 52-89) | |
| **Stage** | | |
| ***NMIBC*** | | |
| **Papilloma** | 1 | 2% |
| **Ta** | 21 | 46% |
| **T1** | 24 | 52% |
| ***MIBC*** | | |
| **T2** | 34 | 82% |
| **T3** | 4 | 9% |
| **T4** | 4 | 9% |
| **Grade** | | |
| **Papilloma** | 1 | 2% |
| **PUNLMP** | 1 | 2% |
| **Low grade** | 28 | 61% |
| **High grade** | 15 | 33% |
| **Unknown** | 1 | 2% |
| **Recurrence** | | |
| **No** | 18 | 39% |
| **Yes** | 18 | 39% |
| **Unknown** | 10 | 22% |
| **Progression** | | |
| **No** | 15 | 33% |
| **Yes** | 11 | 24% |
| **Unknown** | 20 | 43% |
| **Metastasis** | | |
| **No** | 31 | 74% |
| **Yes** | 7 | 17% |
| **Unknown** | 4 | 9% |

**Supplementary Table 2. Antibodies used in this study.**

| | Clone | Isotype | Dilution | Conjugation | Company | Catalog number |
|---|---|---|---|---|---|---|
| **Immunohistochemistry** | | | | | | |
| ***Primary antibodies*** | | | | | | |
| **CD44s Pan Specific antibody** | 2C5 | Mouse IgG | 1:1000 | - | R&D Systems | BBA10 |
| **CD44v6 antibody** | VFF-18 | Mouse IgG | 1:1000 | - | Merck | MAB4073 |
| ***Secondary antibodies*** | | | | | | |
| **Mouse IgG** | - | Donkey IgG | 1:1000 | Biotin | Jackson ImmunoResearch | 715-065-151 |
| **Immunofluorescence** | | | | | | |
| ***Primary antibodies*** | | | | | | |
| **CD44 antibody** | Hermes-1 | Rat IgG | 1:150 | - | ThermoFisher Scientific | MA4400 |
| **CD44v6 antibody** | MA54 | Mouse IgG | 1:250 | - | ThermoFisher Scientific | 33-6700 |
| **E-Cadherin antibody** | 24E10 | Rabbit IgG | 1:400 | - | Cell Signaling Technology | 3195 |
| ***Secondary antibodies*** | | | | | | |
| **Rat IgG** | - | Goat IgG | 1:1000 | Alexa Fluor 647 | ThermoFisher Scientific | A-21247 |
| **Mouse IgG** | - | Goat IgG | 1:1000 | Alexa Fluor 647 | ThermoFisher Scientific | A-21235 |
| **Rabbit IgG** | - | Goat IgG | 1:1000 | Alexa Fluor 488 | ThermoFisher Scientific | A-11008 |
| | **Clone** | **Isotype** | **Final Concentration** | **Fluorochrome** | **Company** | **Catalog number** |
| **Flow cytometry** | | | | | | |
| **CD44 antibody** | J.173 | Mouse IgG | 2 ng/µl (1x$10^5$ cells) | FITC | Beckman Coulter | IM1219U |
| **CD44v6 antibody** | 2F10 | Mouse IgG | 1 ng/µl (1x$10^5$ cells) | PE | BD Biosciences | 566803 |
| **CD326 (EpCAM) antibody** | 9C4 | Mouse IgG | 0.6 ng/µl (1x$10^5$ cells) | APC | BioLegend | 324208 |
| **CD45 antibody** | 30-F11 | Rat IgG | 1.5 ng/µl (1x$10^5$ cells) | PE-Cy7 | eBioscience | 25-0451-82 |

**Supplementary Table 3. Cell lines used in this study.**

| Name | Source | Stage | Grade | Sex | Classification | References |
|---|---|---|---|---|---|---|
| **SW-780** | Transitional Cell Carcinoma | nr | G1 | F | Luminal | 1 |
| **MGH-U3** | Focal severe urothelial atypia | Ta/T1 | G1 | M | Urobasal A/B | 2 |
| **MGH-U4** | Focal severe urothelial atypia | nr | G1 | M | Urobasal A/B | 2 |
| **RT-4** | Transitional Cell Papilloma | T1 | G1-2 | M | Luminal | 3 |
| **5637** | Urothelial Cell Carcinoma | nr | G2 | M | Genomically Unstable Squamous Cell Carcinoma-like | 4-6 |
| **RT-112** | Transitional Cell Carcinoma | Ta | G2 | F | Urobasal A/B | 6-7 |
| **T24** | Transitional Cell Carcinoma | Ta | G3 | F | - | 8 |
| **UM-UC-3** | Urothelial Cell Carcinoma | T2-T4 | nr | M | Basal | 9 |
| **J82** | Transitional Cell Carcinoma | T3 | G3 | M | Genomically Unstable Squamous Cell Carcinoma-like | 10 |
| **253J** | Urothelial Cell Carcinoma | T4 | G4 | M | Basal | 11 |

nr = not reported.

**Supplementary References**

1. Kyriazis AA, Kyriazis AP, McCombs WB 3rd, Peterson WD Jr. Morphological, biological, and biochemical characteristics of human bladder transitional cell carcinomas grown in tissue culture and in nude mice. Cancer Res. 1984 Sep;44(9):3997-4005.
2. Lin CW, Lin JC, Prout GR, Jr.: Establishment and characterization of four human bladder tumor cell lines and sublines with different degrees of malignancy. Cancer research 1985, 45(10):5070-5079.
3. Rigby CC, Franks LM. A human tissue culture cell line from a transitional cell tumour of the urinary bladder: growth, chromosone pattern and ultrastructure. Br J Cancer. 1970 Dec;24(4):746-54.
4. Fogh J, Fogh JM, Orfeo T: One hundred and twenty-seven cultured human tumor cell lines producing tumors in nude mice. Journal of the National Cancer Institute 1977, 59(1):221-226.
5. Rieger KM, Little AF, Swart JM, Kastrinakis WV, Fitzgerald JM, Hess DT, Libertino JA, Summerhayes IC: Human bladder carcinoma cell lines as indicators of oncogenic change relevant to urothelial neoplastic progression. British journal of cancer 1995, 72(3):683-690.
6. Fanning P, Bulovas K, Saini KS, Libertino JA, Joyce AD, Summerhayes IC: Elevated expression of pp60c-src in low grade human bladder carcinoma. Cancer research 1992, 52(6):1457-1462.
7. Masters JR, Hepburn PJ, Walker L, Highman WJ, Trejdosiewicz LK, Povey S, Parkar M, Hill BT, Riddle PR, Franks LM: Tissue culture model of transitional cell carcinoma: characterization of twenty-two human urothelial cell lines. Cancer research 1986, 46(7):3630-3636.
8. Bubeník J, Baresová M, Viklický V, Jakoubková J, Sainerová H, Donner J. Established cell line of urinary bladder carcinoma (T24) containing tumour-specific antigen. Int J Cancer. 1973 May;11(3):765-73.
9. Grossman HB, Wedemeyer G, Ren L, Wilson GN, Cox B. Improved growth of human urothelial carcinoma cell cultures. J Urol. 1986 Oct;136(4):953-9. doi: 10.1016/s0022-5347(17)45139-1.
10. Fogh J: Cultivation, characterization, and identification of human tumor cells with emphasis on kidney, testis, and bladder tumors. National Cancer Institute monograph 1978(49):5-9.
11. Elliott AY, Cleveland P, Cervenka J, Castro AE, Stein N, Hakala TR, Fraley EE. Characterization of a cell line from human transitional cell cancer of the urinary tract. J Natl Cancer Inst. 1974 Nov;53(5):1341-9.

**Supplementary Table 4. Sequence of the oligonucleotides used in this study for reverse transcription and PCR amplification of specific genes.**

| Primers | Sequence |
|---|---|
| **CD44s-F** | 5´-GGAGCAGCACTTCAGGAGGTTAC-3´ |
| **CD44s-R** | 5´-TGTCTTCGTCTGGGATGGGG-3´ |
| **CD44v6-F** | 5´-CCAGGCAACTCCTAGTAGTACAACG-3´ |
| **CD44v6-R** | 5´-CGAATGGGAGTCTTCTTTGGGT-3´ |
| **TBP-F** | 5´-ACAACAGCCTGCCACCTTAC-3´ |
| **TBP-R** | 5´-GCCATAAGGCATCATTGGAC-3´ |
| **GUSB-F** | 5´-CCTGTGACCTTTGTGAGCAA-3´ |
| **GUSB-R** | 5´-AACAGATCACATCCACATACGG-3´ |
| **CDH1-RT** | 5´-TGCTTAACCCCTCACCTTGAAGG-3´ |
| **CDH1-F** | 5´-GGTCTGTCATGGAAGGTGCT-3´ |
| **CDH1-R** | 5´-GATGGCGGCATTGTAGGT-3´ |

#### Supplementary Figures

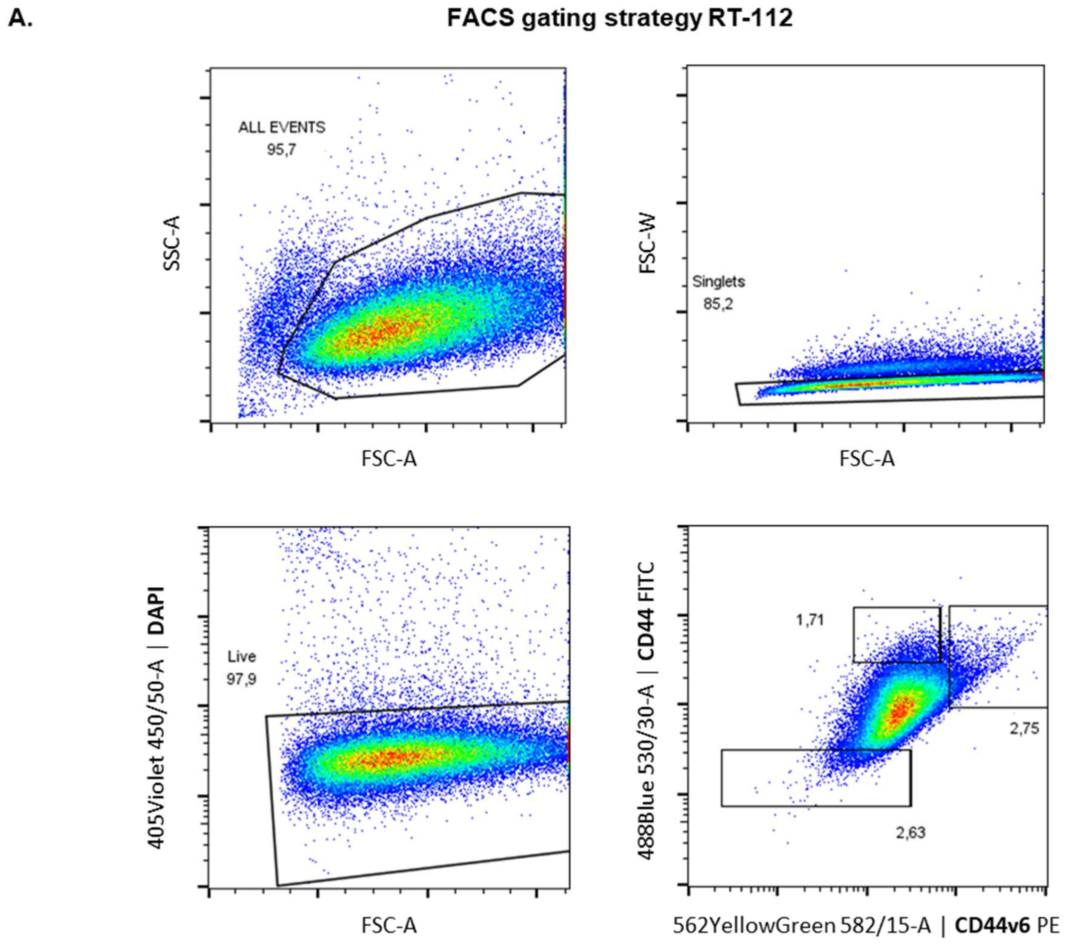

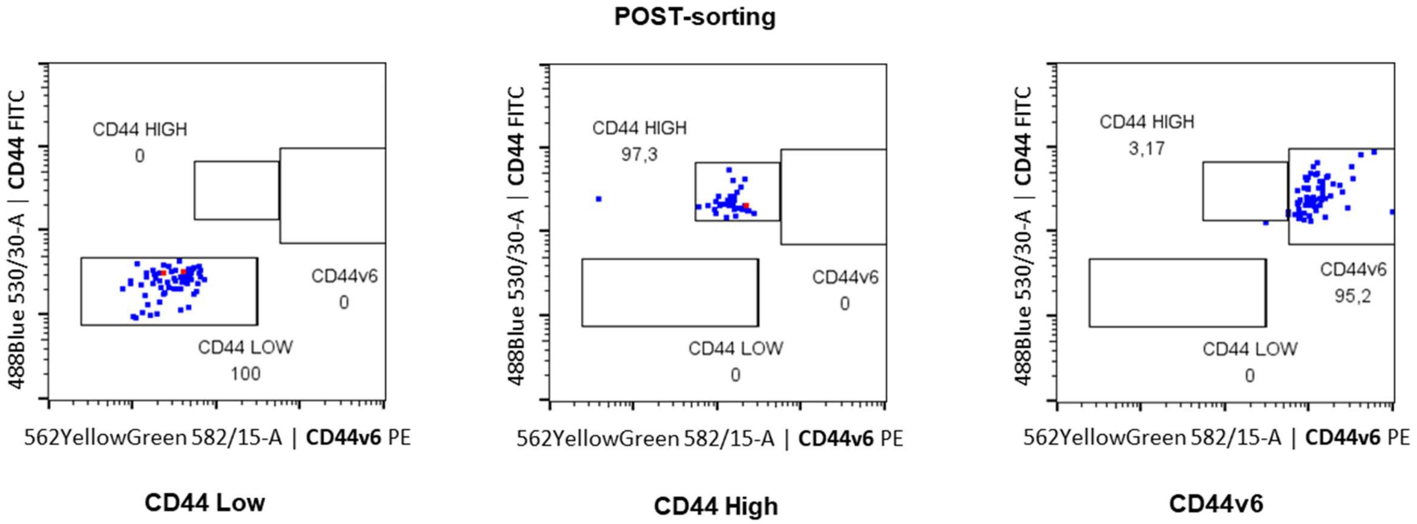

**B.** **FACS gating strategy J82**

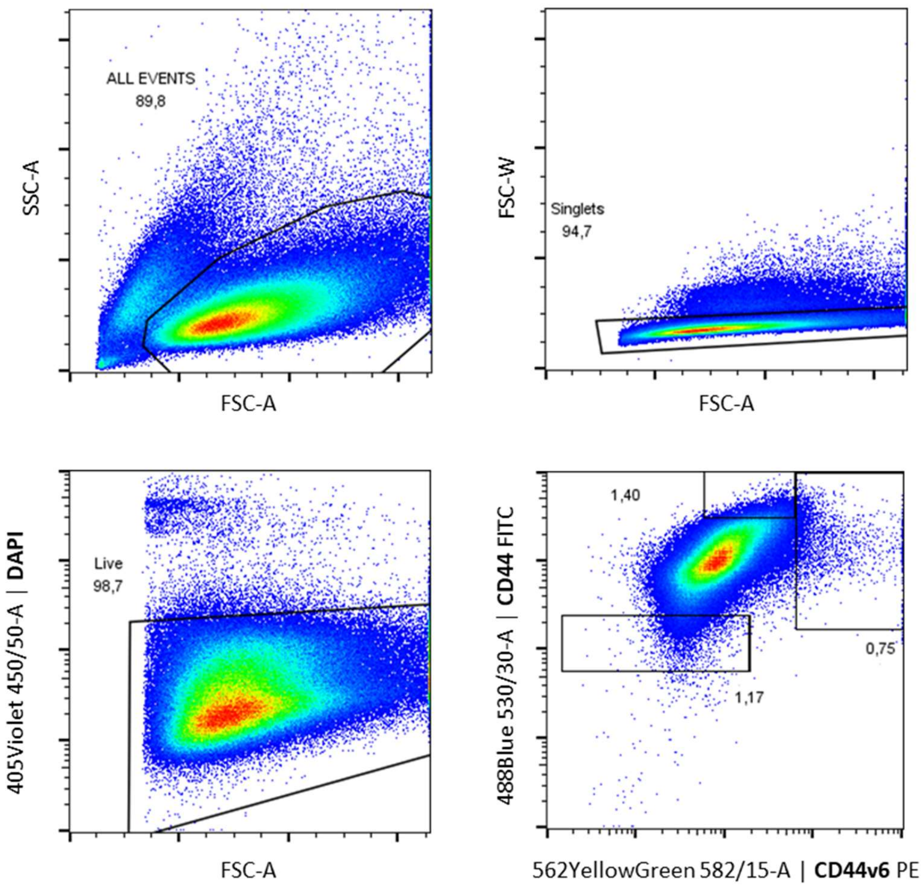

**POST-sorting**

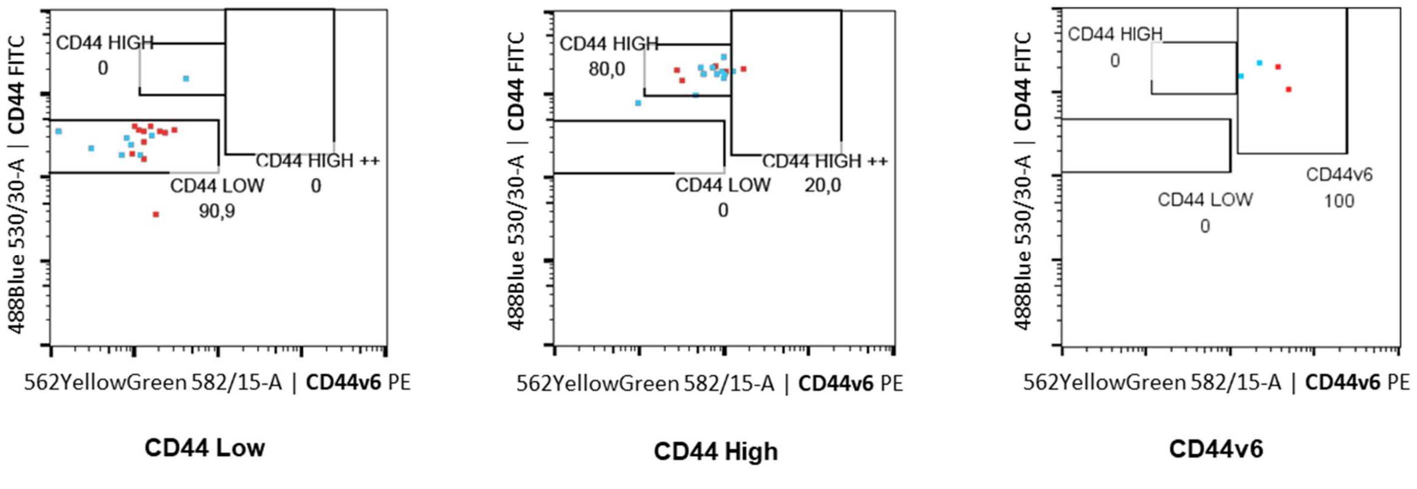

**Supplementary Figure 1. FACS gating strategy used to isolate CD44 Low, CD44 High and CD44v6 High cell populations from RT-112 (A) and J82 (B) parental cell lines.**

A.

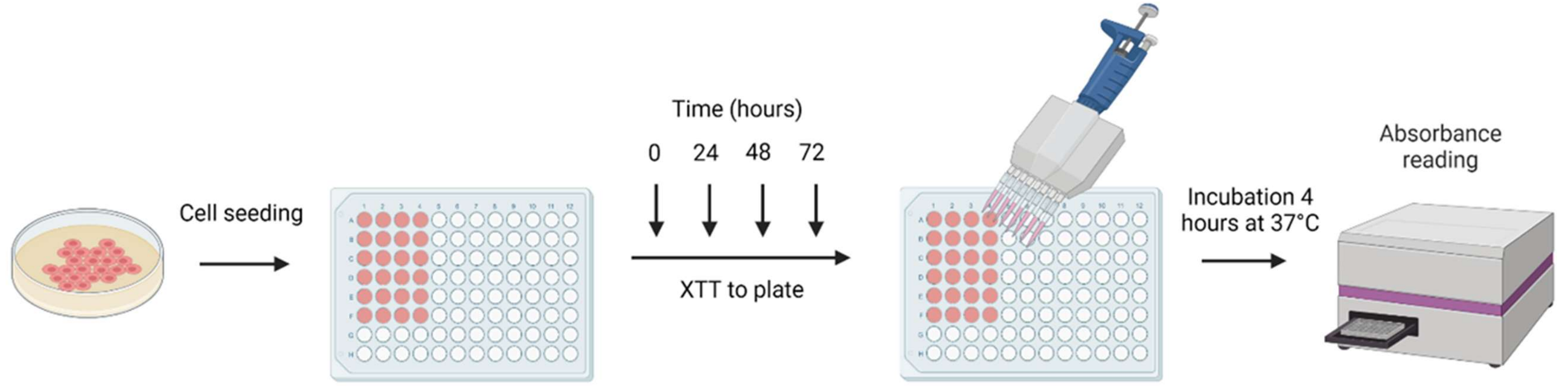

B.

**RT-112**

| | 48h | 72h |
|---|---|---|
| Parental vs. CD44 Low | ns | ** |
| Parental vs. CD44 High | * | *** |
| Parental vs. CD44v6 | ns | ns |
| CD44 Low vs. CD44 High | *** | **** |
| CD44 Low vs. CD44v6 | ns | ** |
| CD44 High vs. CD44v6 | ns | *** |

**J82**

| | 48h | 72h |
|---|---|---|
| Parental vs. CD44 Low | ns | * |
| Parental vs. CD44 High | ns | ** |
| Parental vs. CD44v6 | ns | **** |
| CD44 Low vs. CD44 High | ns | **** |
| CD44 Low vs. CD44v6 | * | **** |
| CD44 High vs. CD44v6 | ns | ns |

**Supplementary Figure 2. Evaluation of cell proliferation *in vitro*.** A) Schematic representation of the proliferation assay with the XTT Cell proliferation kit. B) Statistical significance tables for the comparisons of proliferative capacity between the parental cell lines (RT-112 and J82) and their respective CD44 Low, CD44 High and CD44v6 cell lines. ns = not significant, *p-value<0.05, **p-value<0.01, ***p-value<0.001, ****p-value<0.0001.

A.

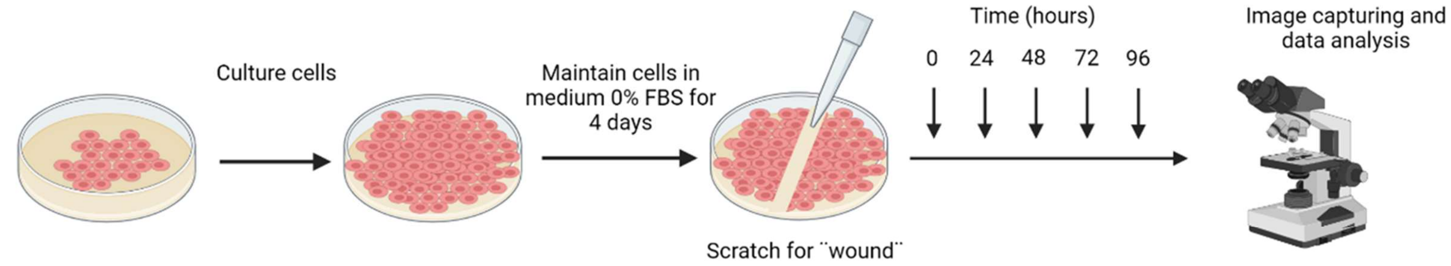

B.

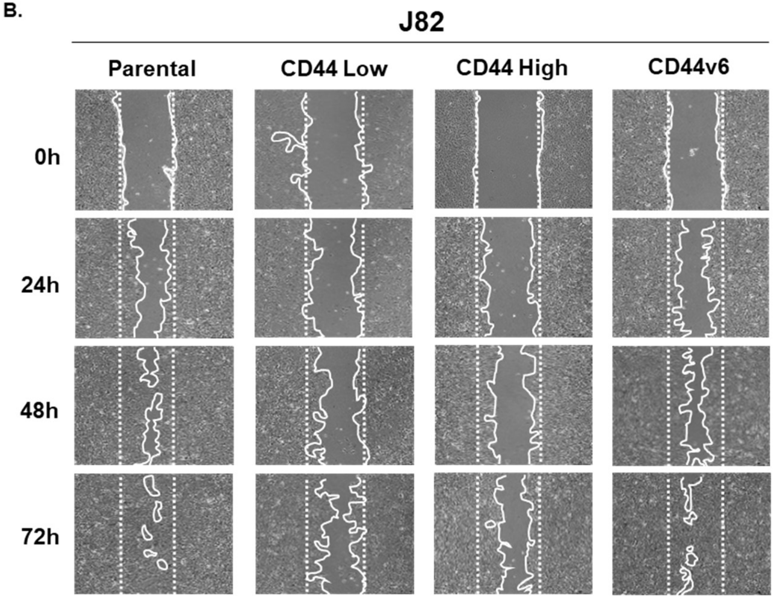

C.

**RT-112**

| | 24h | 48h | 72h |
|---|---|---|---|
| Parental vs. CD44 Low | ns | ** | *** |
| Parental vs. CD44 High | **** | **** | **** |
| Parental vs. CD44v6 | ** | * | ** |
| CD44 Low vs. CD44 High | **** | **** | **** |
| CD44 Low vs. CD44v6 | ns | ns | ns |
| CD44 High vs. CD44v6 | **** | **** | **** |

**J82**

| | 24h | 48h | 72h |
|---|---|---|---|
| Parental vs. CD44 Low | ns | ns | ns |
| Parental vs. CD44 High | ns | ns | ** |
| Parental vs. CD44v6 | **** | ns | ns |
| CD44 Low vs. CD44 High | ns | ns | ns |
| CD44 Low vs. CD44v6 | *** | ns | ns |
| CD44 High vs. CD44v6 | *** | ns | ** |

**Supplementary Figure 3. Evaluation of cell migration *in vitro*.** A) Schematic representation of the Wound Scratch Assay. B) Representative image of J82 wound healing after 24 h, 48 h and 72 h of wound scratching. C) Statistical significance tables for the comparison of migration capacity between the parental cell lines (RT-112 and J82) and their respective CD44 Low, CD44 High and CD44v6 cell lines after 24 h, 48 h and 72 h of wound scratching. ns = not significant, *p-value<0.05, **p-value<0.01, ***p-value<0.001, ****p-value<0.0001.

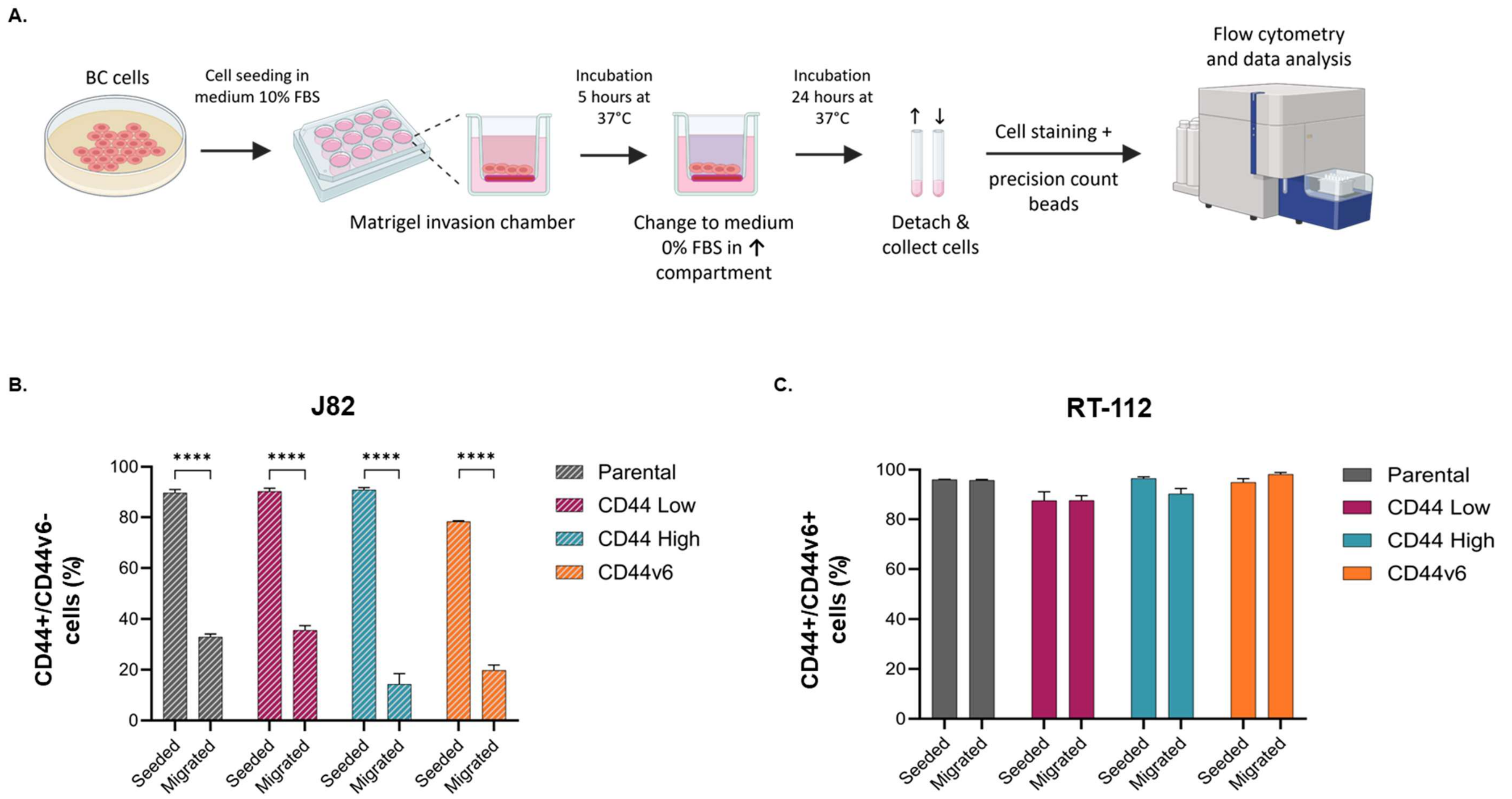

**Supplementary Figure 4. Evaluation of cell invasion *in vitro*.** A) Schematic representation of the cell invasion assay, using Boyden Chambers. B) Percentage of CD44+/CD44v6- cells among the seeded and invading J82 cell populations. C) Percentage of CD44+/CD44v6+ cells among the seeded and invading RT-112 cell populations. Error bars represent the mean ± SEM. *p-value<0.05, **p-value<0.01, ****p-value<0.0001.

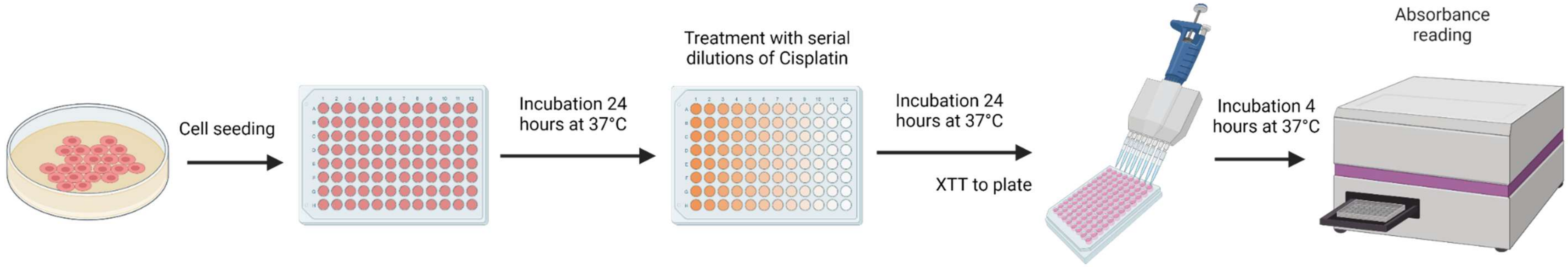

**Supplementary Figure 5. Evaluation of cisplatin sensitivity *in vitro*.** Schematic representation of the viability assay with the XTT Cell proliferation kit.

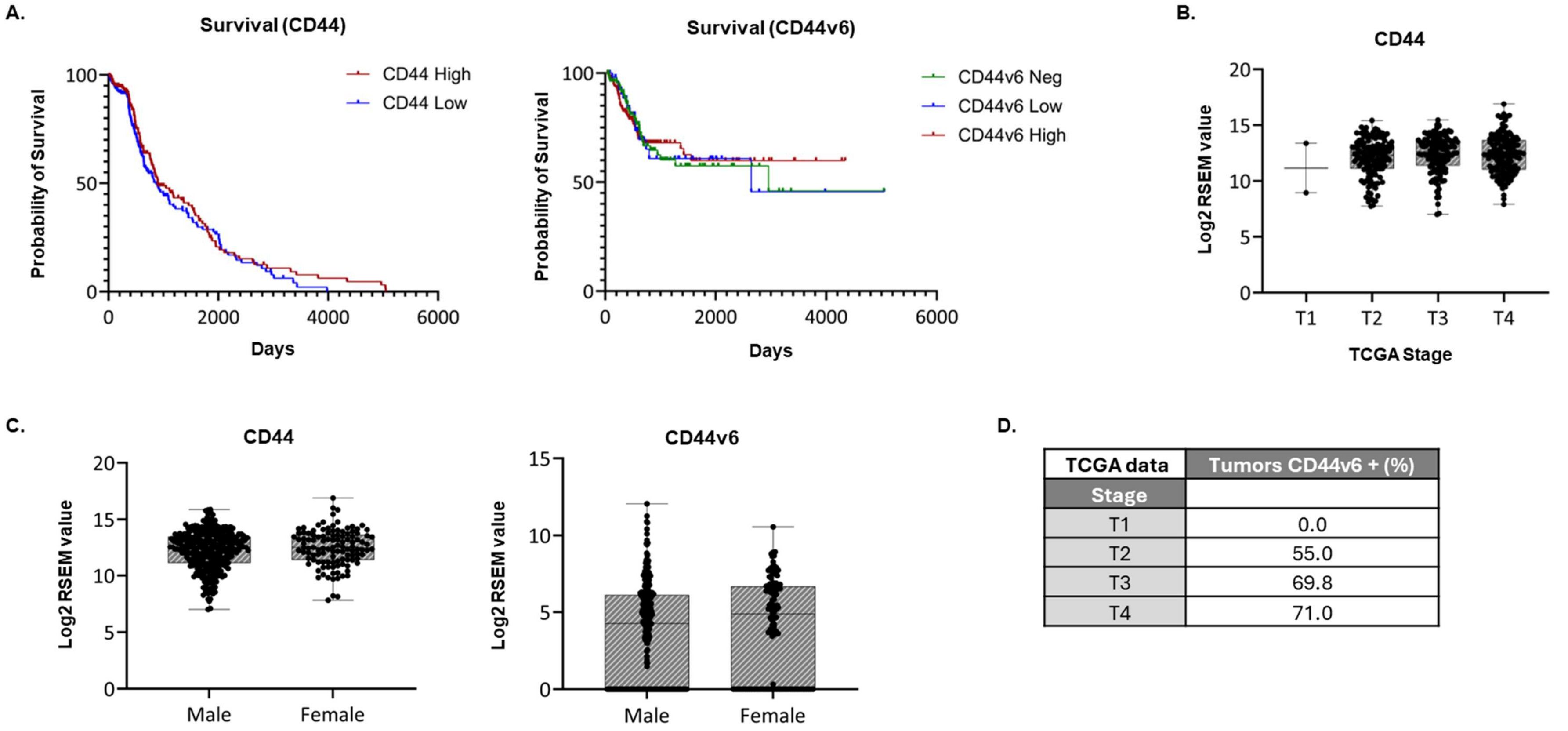

| TCGA data | Tumors CD44v6 + (%) |
|---|---|
| Stage | |
| T1 | 0.0 |
| T2 | 55.0 |
| T3 | 69.8 |
| T4 | 71.0 |

**Supplementary Figure 6. Analysis of *CD44* and *CD44v6* gene expression in BC patient samples.** Gene expression data were obtained from the TCGA database using the TSVdb web-tool (Supplementary Table 1). A) Kaplan–Meier curves representing the overall survival of BC patients based on CD44 and CD44v6 gene expression levels. B) Evaluation of relative *CD44s* gene expression levels across 406 BC patients based on tumor stage. Patients from stage I to stage IV disease were included in this analysis. C) Relative CD44s and CD44v6 gene expression analysis between BC patient samples based on sex. D) Table representing the percentage (%) of CD44v6-positive (CD44v6+) tumors for each disease stage, based on CD44v6 gene expression data. Boxplots were used for representation, showing all individual values. *p-value<0.05.

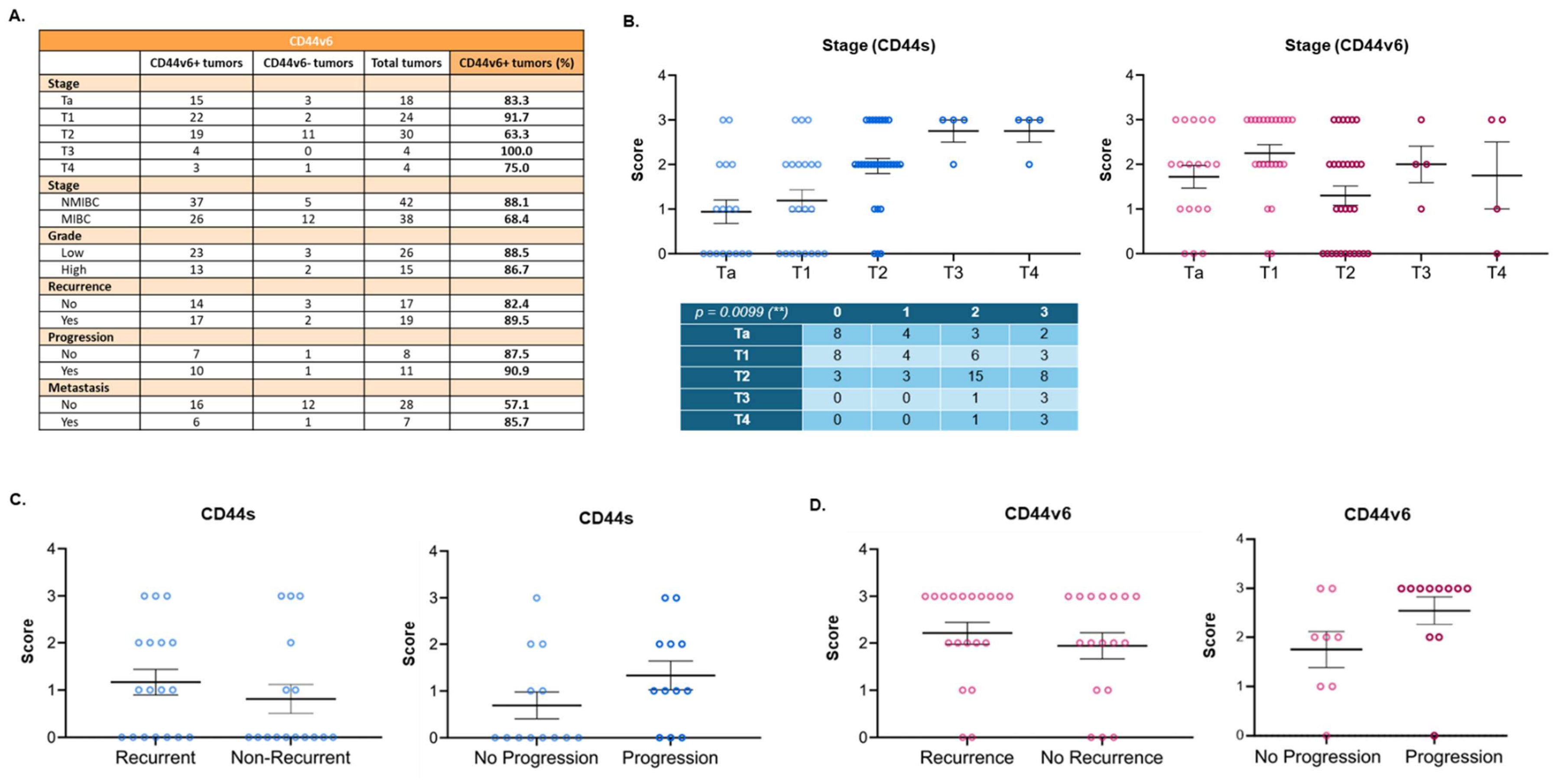

A.

| CD44v6 | | | | |
|---|---|---|---|---|
| | CD44v6+ tumors | CD44v6- tumors | Total tumors | CD44v6+ tumors (%) |
| **Stage** | | | | |
| Ta | 15 | 3 | 18 | 83.3 |
| T1 | 22 | 2 | 24 | 91.7 |
| T2 | 19 | 11 | 30 | 63.3 |
| T3 | 4 | 0 | 4 | 100.0 |
| T4 | 3 | 1 | 4 | 75.0 |
| **Stage** | | | | |
| NMIBC | 37 | 5 | 42 | 88.1 |
| MIBC | 26 | 12 | 38 | 68.4 |
| **Grade** | | | | |
| Low | 23 | 3 | 26 | 88.5 |
| High | 13 | 2 | 15 | 86.7 |
| **Recurrence** | | | | |
| No | 14 | 3 | 17 | 82.4 |
| Yes | 17 | 2 | 19 | 89.5 |
| **Progression** | | | | |
| No | 7 | 1 | 8 | 87.5 |
| Yes | 10 | 1 | 11 | 90.9 |
| **Metastasis** | | | | |
| No | 16 | 12 | 28 | 57.1 |
| Yes | 6 | 1 | 7 | 85.7 |

B.

| *p = 0.0099 (**)* | 0 | 1 | 2 | 3 |
|---|---|---|---|---|
| Ta | 8 | 4 | 3 | 2 |
| T1 | 8 | 4 | 6 | 3 |
| T2 | 3 | 3 | 15 | 8 |
| T3 | 0 | 0 | 1 | 3 |
| T4 | 0 | 0 | 1 | 3 |

**Supplementary Figure 7. CD44 and CD44v6 protein expression analysis using BC patient samples.** A) Table representing the percentage (%) of CD44v6-positive (CD44v6+) tumors for each clinical factor evaluated in this study. Progression is defined as the appearance of MIBC in an initial NMIBC patient. B) Evaluation of CD44s and CD44v6 receptor expression among BC patient samples based on tumor stage. Patients from stage Ta to stage IV disease were included in this analysis. C) Analysis of CD44s receptor expression between BC patient samples based on recurrence-free survival and disease progression. D) Evaluation of CD44v6 receptor expression among BC patient samples based on recurrence-free survival and disease progression. Error bars represent the mean ± SEM. **p-value<0.01.

A.

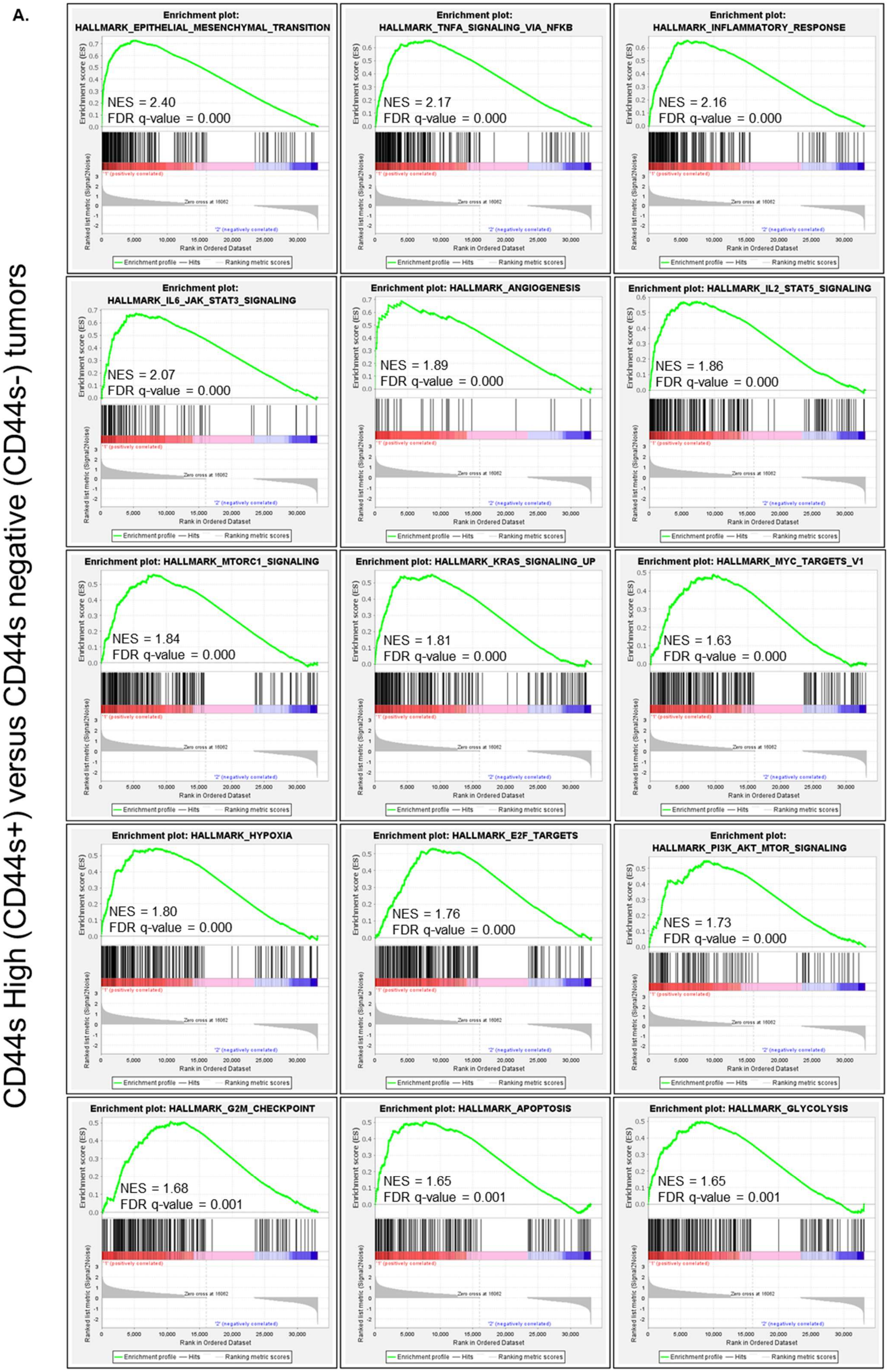
CD44s High (CD44s+) versus CD44s negative (CD44s-) tumors
Enrichment plot: HALLMARK_EPITHELIAL_MESENCHYMAL_TRANSITION
NES = 2.40
FDR q-value = 0.000
Enrichment plot: HALLMARK_TNFA_SIGNALING_VIA_NFKB
NES = 2.17
FDR q-value = 0.000
Enrichment plot: HALLMARK_INFLAMMATORY_RESPONSE
NES = 2.16
FDR q-value = 0.000
Enrichment plot: HALLMARK_IL6_JAK_STAT3_SIGNALING
NES = 2.07
FDR q-value = 0.000
Enrichment plot: HALLMARK_ANGIOGENESIS
NES = 1.89
FDR q-value = 0.000
Enrichment plot: HALLMARK_IL2_STAT5_SIGNALING
NES = 1.86
FDR q-value = 0.000
Enrichment plot: HALLMARK_MTORC1_SIGNALING
NES = 1.84
FDR q-value = 0.000
Enrichment plot: HALLMARK_KRAS_SIGNALING_UP
NES = 1.81
FDR q-value = 0.000
Enrichment plot: HALLMARK_MYC_TARGETS_V1
NES = 1.63
FDR q-value = 0.000
Enrichment plot: HALLMARK_HYPOXIA
NES = 1.80
FDR q-value = 0.000
Enrichment plot: HALLMARK_E2F_TARGETS
NES = 1.76
FDR q-value = 0.000
Enrichment plot: HALLMARK_PI3K_AKT_MTOR_SIGNALING
NES = 1.73
FDR q-value = 0.000
Enrichment plot: HALLMARK_G2M_CHECKPOINT
NES = 1.68
FDR q-value = 0.001
Enrichment plot: HALLMARK_APOPTOSIS
NES = 1.65
FDR q-value = 0.001
Enrichment plot: HALLMARK_GLYCOLYSIS
NES = 1.65
FDR q-value = 0.001
Enrichment score (ES)
Ranked list metric (Signal2Noise)
Rank in Ordered Dataset
Zero cross at 16062
'1' (positively correlated)
'2' (negatively correlated)
Enrichment profile — Hits — Ranking metric scores

B.

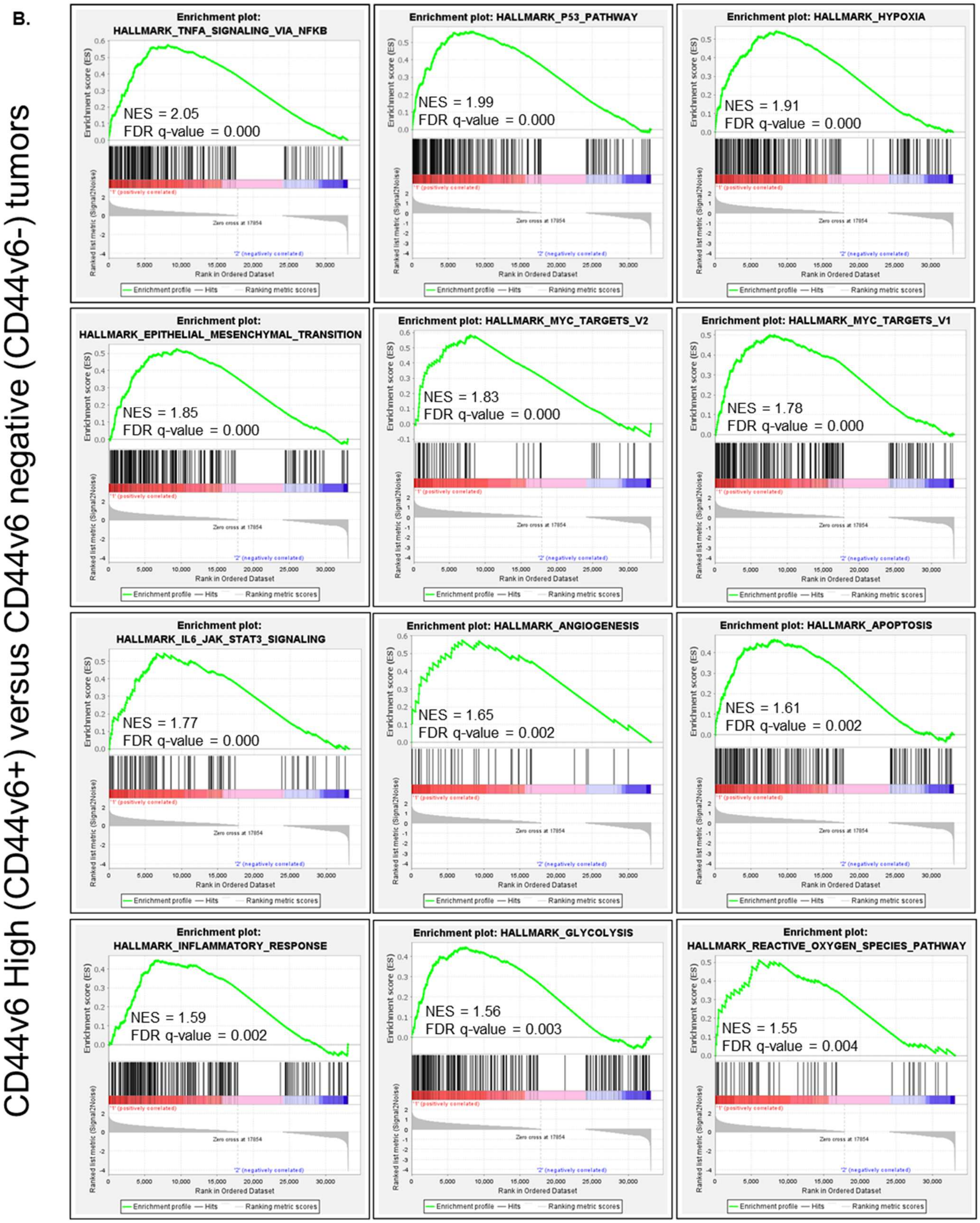

**Supplementary Figure 8. GSEA analysis, comparing A) CD44s High (CD44s+) and CD44s negative (CD44s-) tumors, and B) CD44v6 High (CD44v6+) and CD44v6 negative (CD44v6-) tumors.** Normalized enrichment score (NES) and False discovery rate (FDR) q-value are shown.

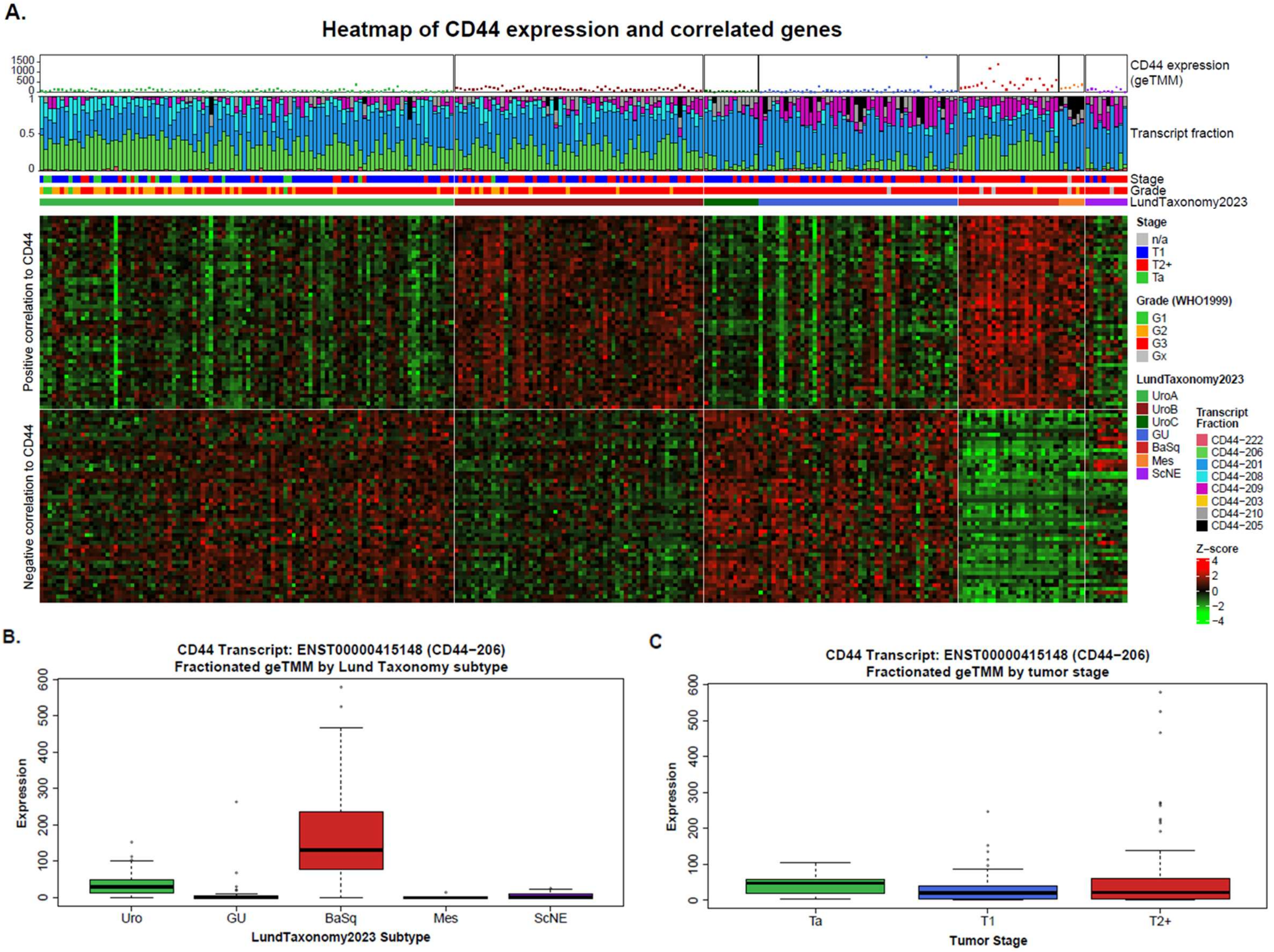

**Supplementary Figure 9. Expression patterns of CD44 and CD44v6 in the Lund "s265" RNA-seq cohort (Cotillas et al., 2024).** A) Panels from top to bottom: Total CD44 expression (geTMM normalized data) across the cohort, individual transcript fractions, stage, grade (WHO1999), Lund Taxonomy subtype classification, expression of CD44 and 49 genes with highest correlation to CD44, expression of 50 inversely correlated genes. B) CD44v6 expression in Lund Taxonomy subtypes. C) CD44v6 expression by tumor stage. Transcript fractions are calculated as each transcripts contribution to total CD44 expression in transcript per million (TPM) format. CD44v6 expression in B and C is calculated as the transcript fraction of ENST00000415148 multiplied with total CD44 expression in cohort-normalized geTMM format.

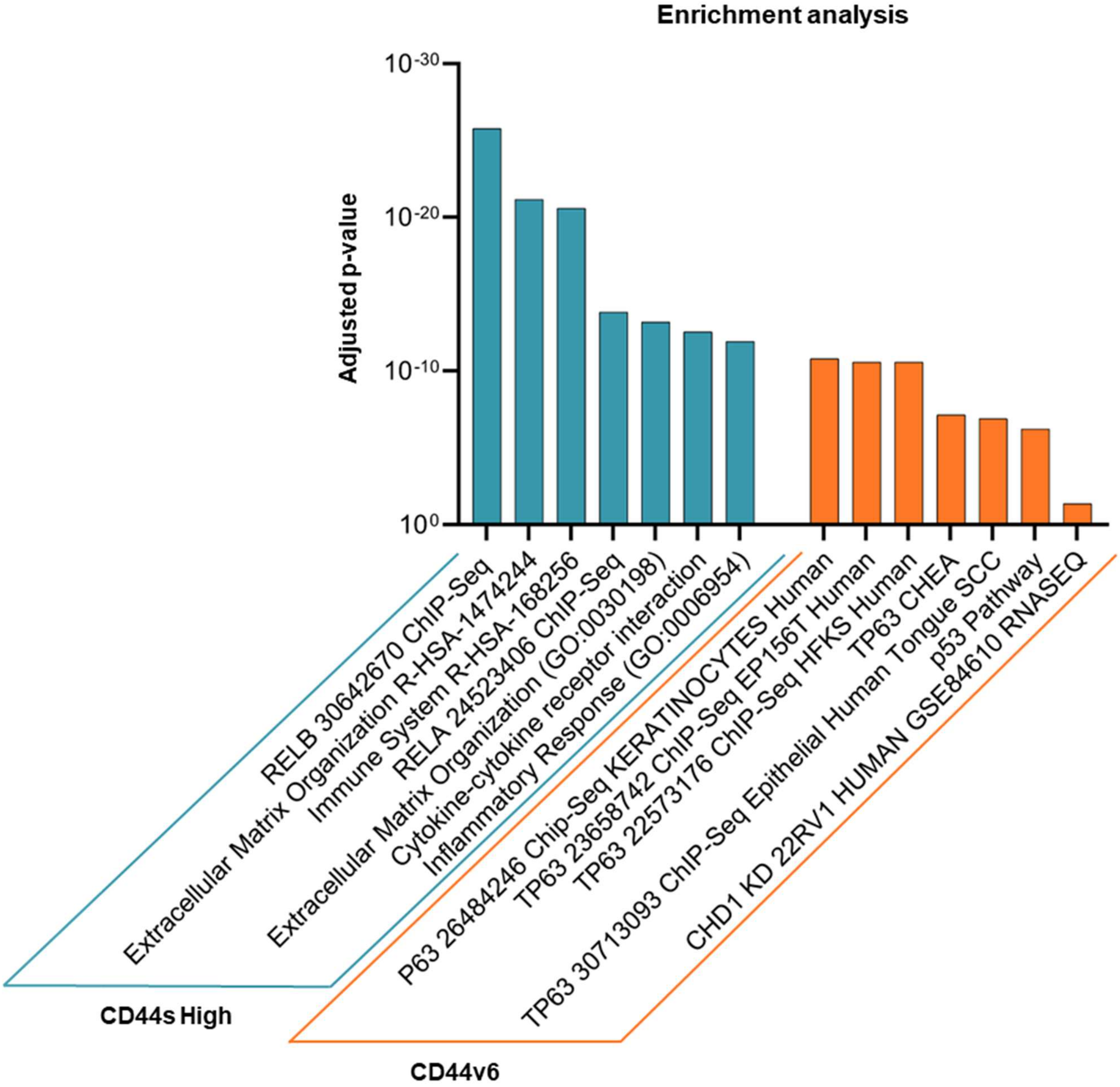

**Supplementary Figure 10. Enrichment analysis based on DE genes between CD44s High and CD44s negative tumors (CD44), as well as CD44v6 High and CD44v6 negative tumors (CD44v6).**

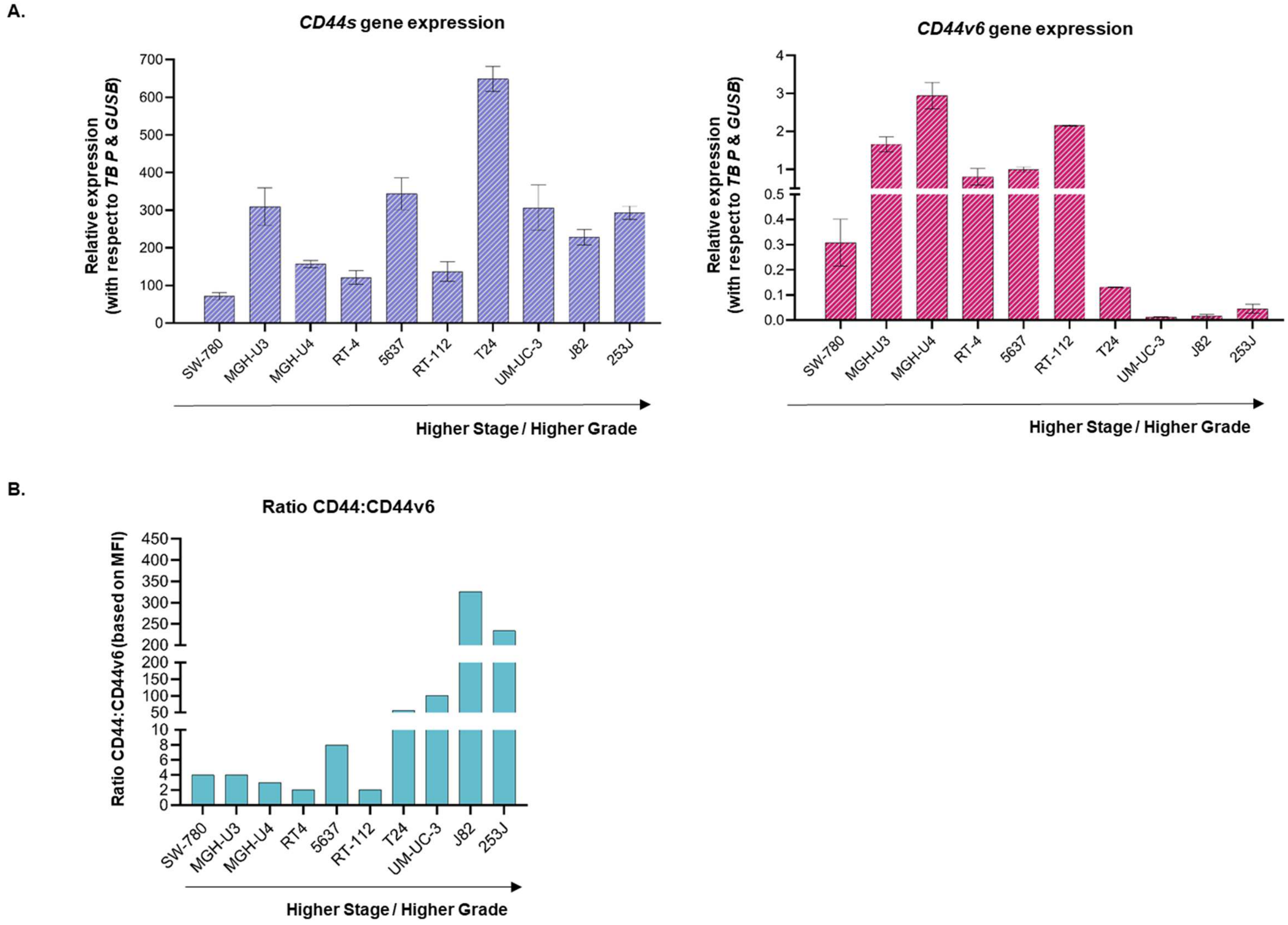

**Supplementary Figure 11. CD44 and CD44v6 expression among human BC cell lines.** A) CD44s and CD44v6 gene expression levels relative to the *TBP* & *GUSB* housekeeping genes. B) CD44:CD44v6 receptor expression ratios, based on MFI. Error bars represent the mean ± SEM.

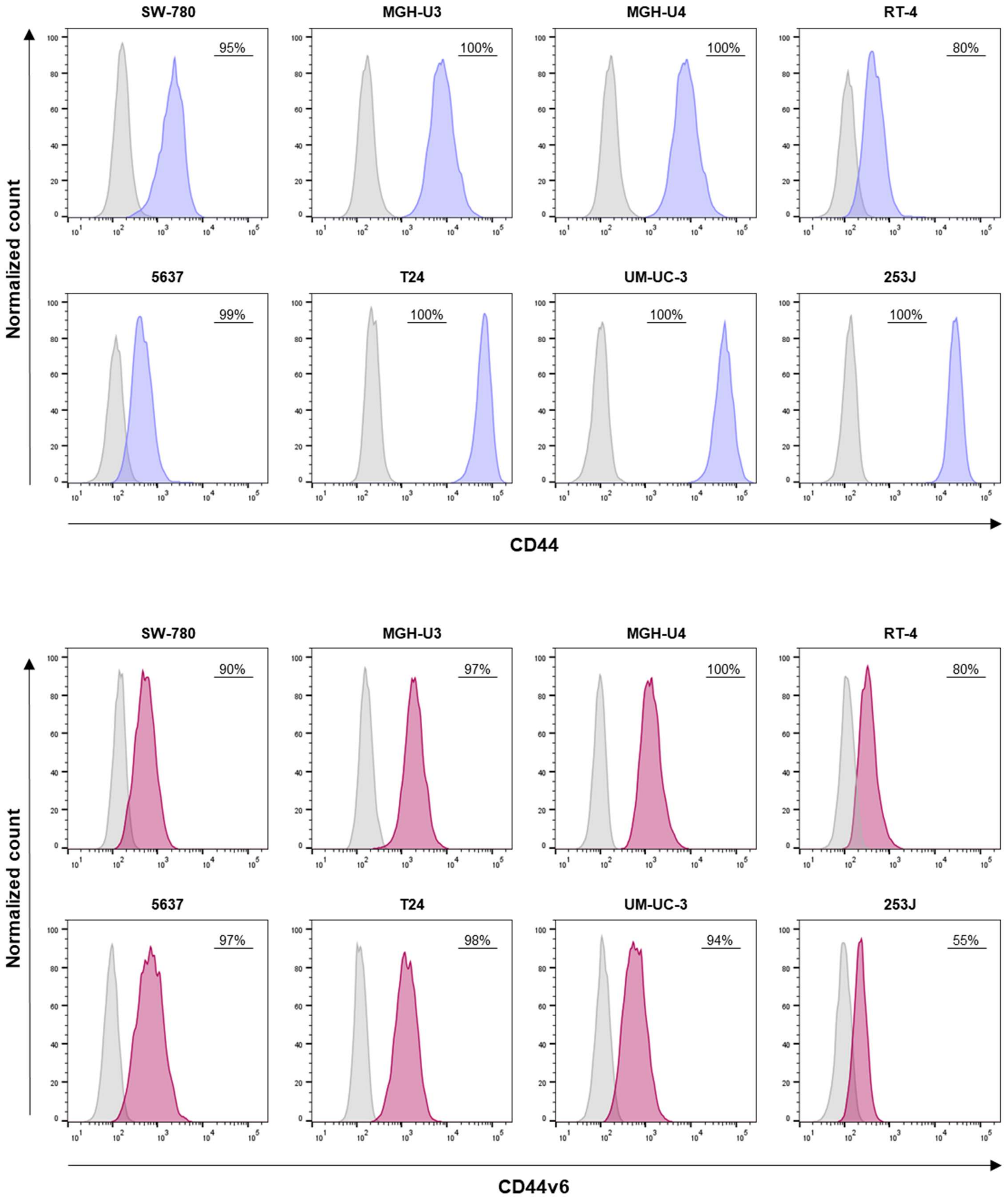

**Supplementary Figure 12. Percentages of CD44+ and CD44v6+ cells across a variety of human BC cell lines.**

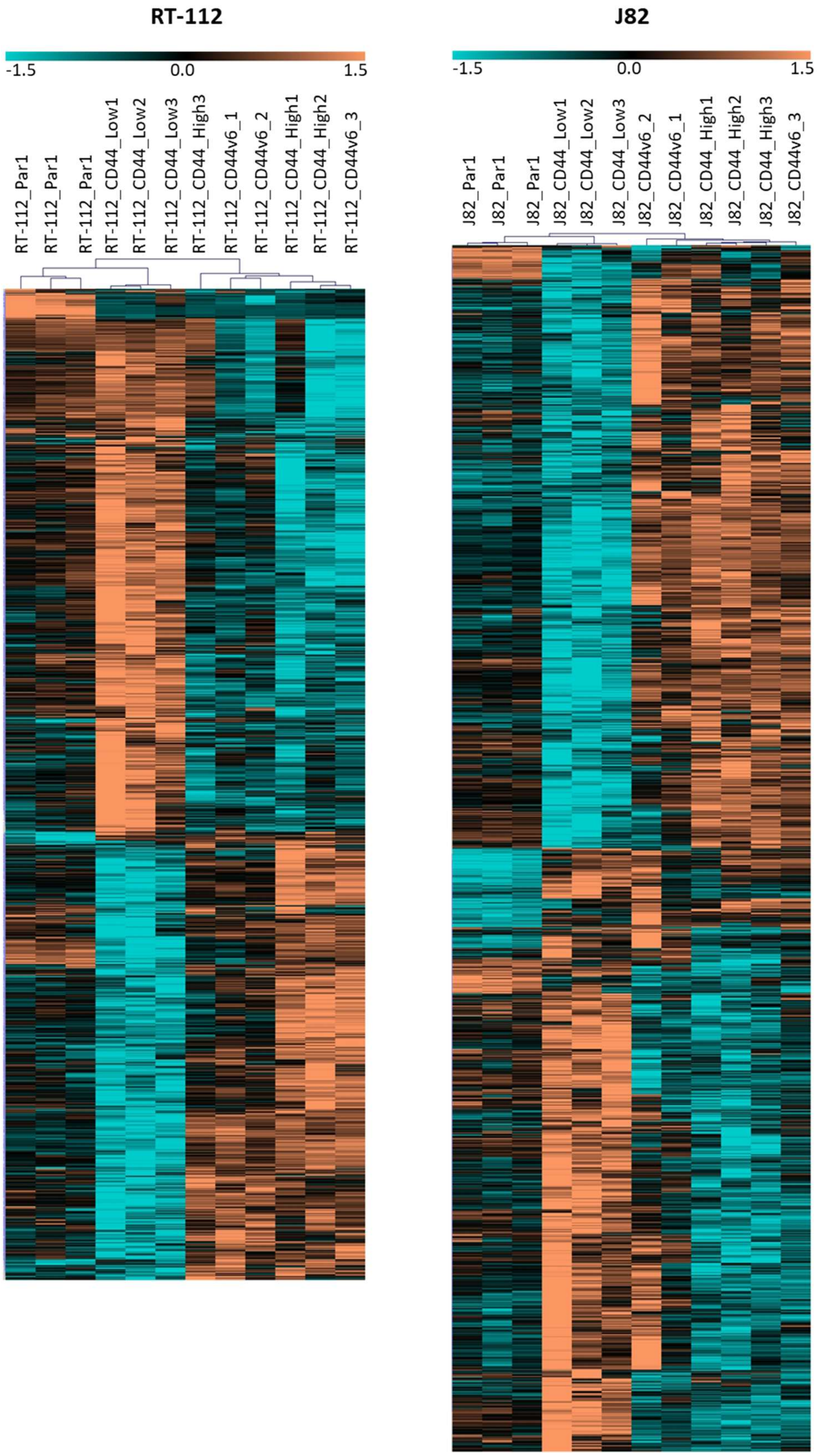

**Supplementary Figure 13. Heatmaps representing unsupervised clustering of the parental cell lines (RT-112 and J82) and their respective newly generated CD44 Low, CD44 High and CD44v6 cell lines, based on gene expression.**

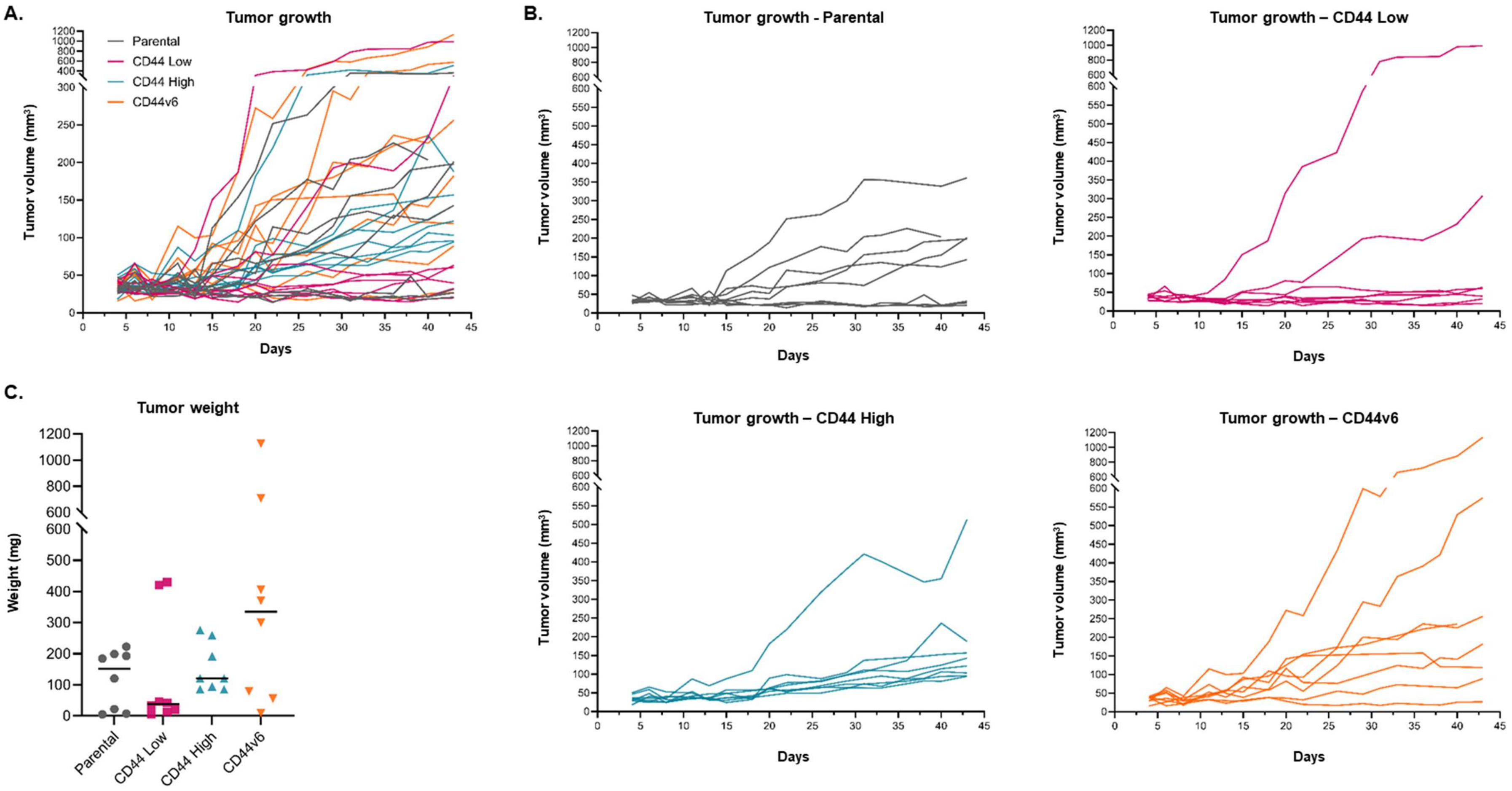

**Supplementary Figure 14. Evaluation of tumorigenesis *in vivo*.** A) Spiderplot representing all individual tumor growth curves after injection of RT-112 parental, CD44 Low, CD44 High and CD44v6 cell lines. B) Spiderplots showing individual tumor growth curves after injection of either RT-112 parental, CD44 Low, CD44 High or CD44v6 cell lines. C) Chart representing the weight of each tumor at the study endpoint. The median values are also shown.

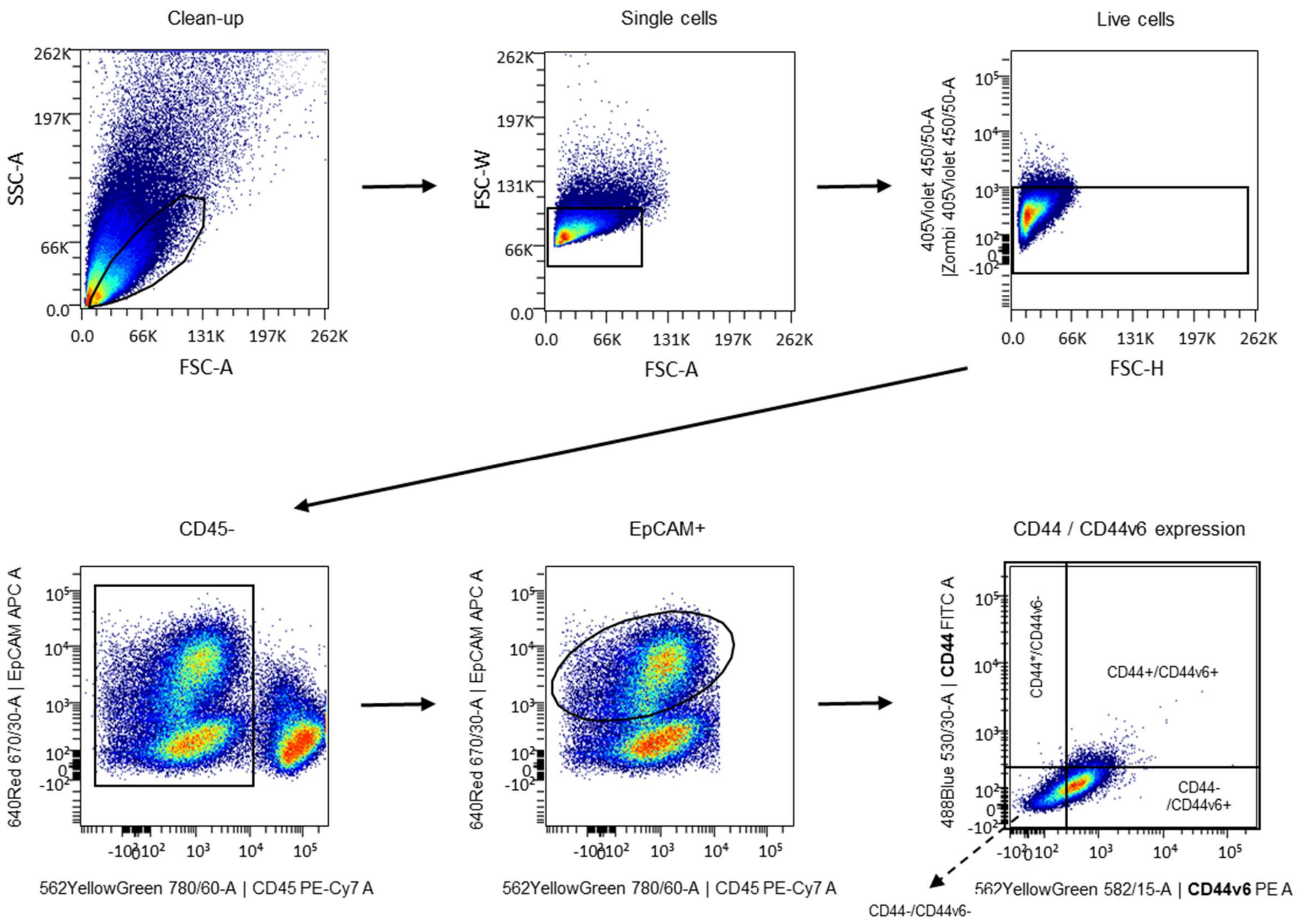

**Supplementary Figure 15. Gating strategy for CD44 and CD44v6 receptor expression analysis among tumors, using flow cytometry and the OMIQ Data Science Platform.** An anti-CD45 antibody was used to discard murine immune cells, and an anti-EpCAM antibody was used to identify tumor cells. Antibodies against CD44 and CD44v6 were used to measure their respective expression levels. The antibodies used in this study are listed in Supplementary Table 2.

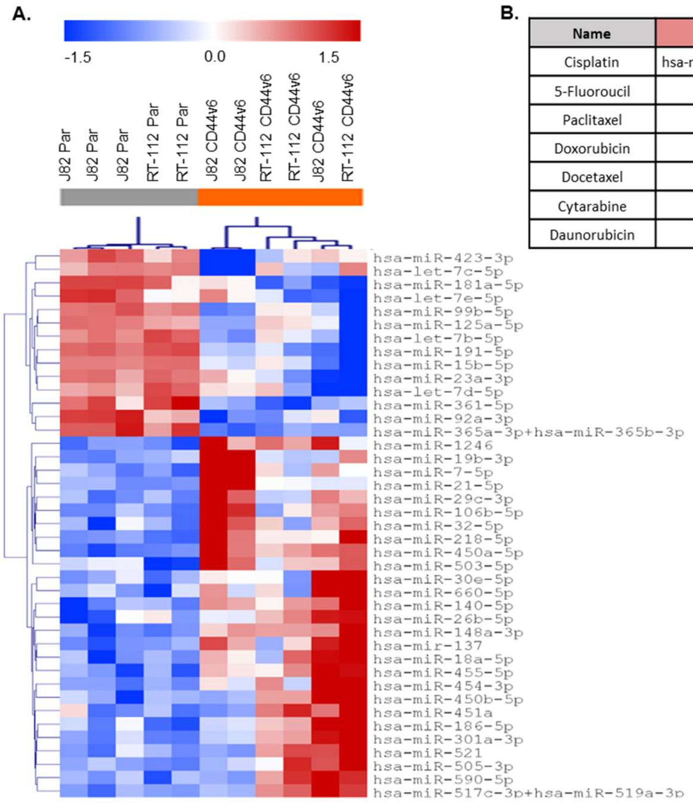

B.

| Name | miRNAs Up | miRNAs Down |
|---|---|---|
| Cisplatin | hsa-miR-15b-5p, hsa-miR-23a-3p | hsa-miR-106b-5p, hsa-miR-21-5p |
| 5-Fluoroucil | | hsa-miR-140-5p, hsa-miR-21-5p |
| Paclitaxel | | hsa-miR-148a-3p, hsa-miR-21-5p |
| Doxorubicin | | hsa-miR-21-5p, hsa-miR-451a |
| Docetaxel | | hsa-miR-21-5p, hsa-miR-505-3p |
| Cytarabine | hsa-miR-181a-5p | hsa-miR-21-5p |
| Daunorubicin | hsa-miR-181a-5p | hsa-miR-21-5p |

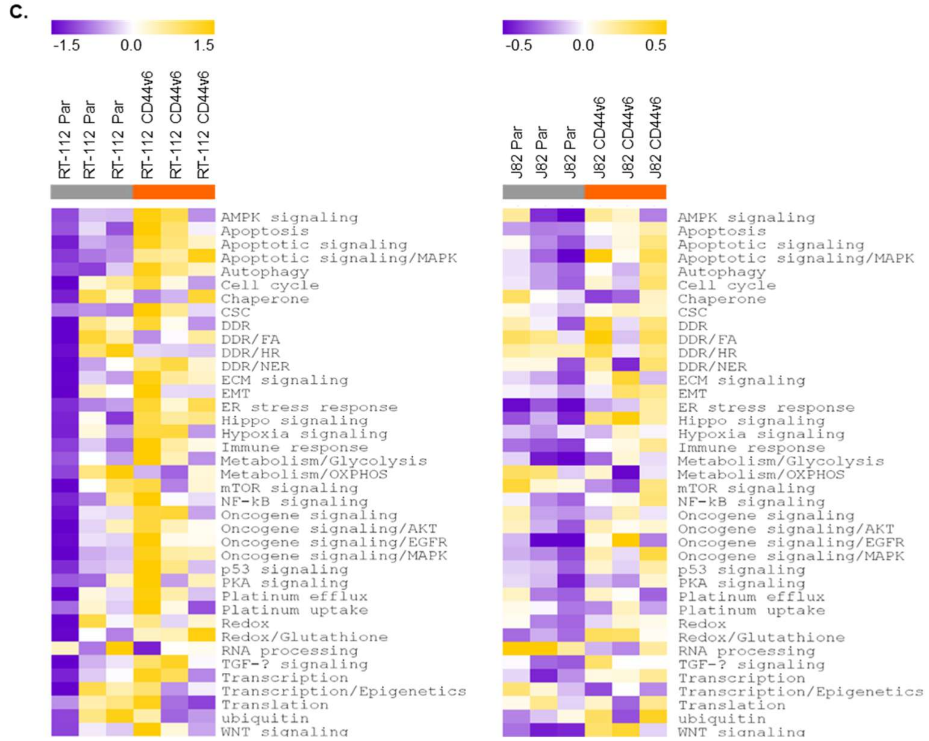

**Supplementary Figure 16. miRNA analysis comparing parental cell lines with generated CD44v6 cell lines.** A) Heatmaps representing unsupervised clustering of DE miRNAs of parental cell lines (RT-112 and J82) and their respective CD44v6 cell lines. B) Table representing DE miRNAs between parental and CD44v6 cell lines, and their association with drug resistance. C) GSVA analysis of DE miRNAs between parental and CD44v6 cell lines and their association with cisplatin resistance-related pathway regulation.
